# Evidence for homing behaviour consistent with path integration in *Dascyllus trimaculatus*

**DOI:** 10.1101/2024.07.22.604582

**Authors:** Freddie Murley, Adelaide Sibeaux, Theresa Burt de Perera

## Abstract

Path integration is a strategy that allows animals to monitor their movement relative to a starting point using directional and distance cues. It has been observed in a wide range of terrestrial species, but evidence of this behaviour in fish is still lacking. In contrast to most animals shown to navigate via path integration, fish are not surface bound but exist within a three-dimensional medium. This may present additional challenges when monitoring their position relative to a specific point. We developed a novel experimental paradigm to test whether the coral reef-dwelling domino damselfish *Dascyllus trimaculatus* could use path integration to navigate back to a shelter location. This consisted of a circular pool with a shelter and landmarks near the edge and a trap containing food in the centre. Our results show that these fish follow homing trajectories back to the previous location of the removed shelter without using landmarks after a foraging trip into the central trap. Error in homing trajectory increased with outward path length, consistent with the use of path integration. These results add *D. trimaculatus* to the growing list of path-integrating species, opening new avenues for investigating the evolution and neural underpinnings of this navigational strategy.

## Introduction

Path integration is a powerful navigational strategy that allows animals to return to a home location efficiently at the end of an outward journey. A path integrating animal must keep track of its directional change and distance travelled on an outward trip using movement relative to direction cues and odometer cues. This is essential to continually update a homing vector, which is the most direct path back to a home location (Müller & Wehner, 1988a; Patel & Cronin, 2020b, Figure 1).

**Figure 1.**
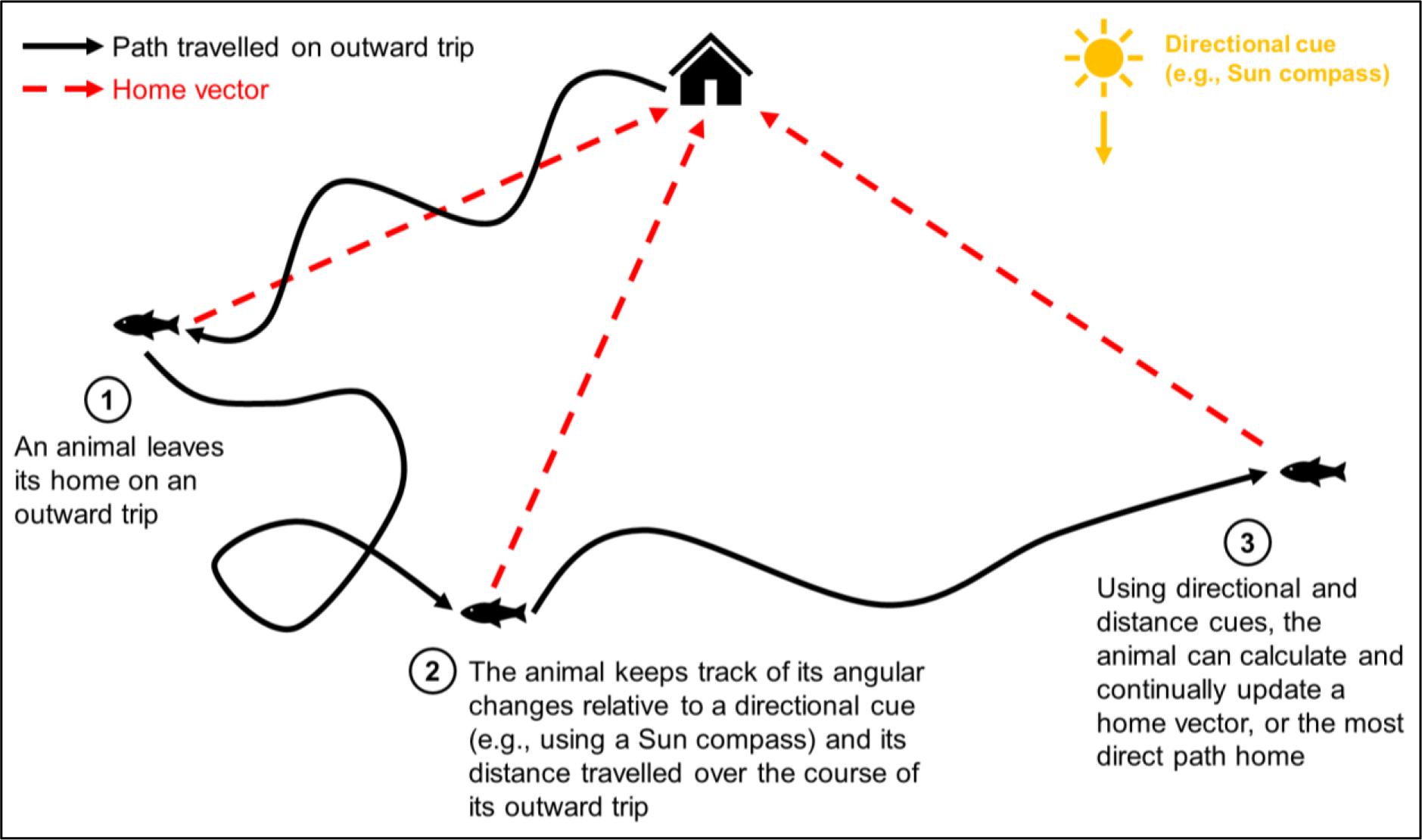
How an animal can keep track of its location relative to its home and calculate the most direct route back via path integration.

The greater the distance a path-integrating animal travels on an outward journey, the more error is predicted to accumulate around its estimate of home location. This is due to the cumulative effect of small errors in directional change and distance measurements over the course of the trip due to inherent sensorimotor system and environmental noise (Cheung, 2014; Cheung et al., 2007). To compensate for the accumulation of errors, many animals perform search behaviour if they don’t reach the exact location of their home using their self-generated homing vector (Patel & Cronin, 2020c). Searching animals often follow ever increasing loops around the end of their homing vector, returning to the start of their search at the end of each loop (Wehner & Srinivasan, 1981). The total journey of a path integrating animal leaving its shelter for a foraging trip can therefore be broken down into (1) an outward trip, (2) a homeward path following a homing vector, and (3) search behaviour if the homing vector is inaccurate.

Animals can use allothetic or idiothetic directional cues for path integration. Allothetic directional cues are derived from the environment and include stable external directional cues like compasses based on the Sun’s position or Earth’s magnetic field. Idiothetic cues are generated from self-motion and can incorporate information from the proprioceptive or vestibular systems. Error accumulates more quickly if using idiothetic cues than if using allothetic cues for path integration (Cheung et al., 2007). The use of idiothetic directional cues requires the cumulative use of rotation estimates, leading to a rapid accumulation of errors. In contrast, the use of allothetic cues allows the constant calibration of an animal’s direction estimate relative to a stable external cue, so directional error should not accumulate. Irrespective of the type of directional cue used, errors around the estimation of distance travelled should always accumulate linearly with increasing outward path length. Therefore, animals using allothetic directional cues to path integrate should exhibit a linear accumulation of error around their estimated home location with increasing outward path length. In contrast, animals using idiothetic cues should exhibit a more rapid accumulation of error with increasing outward path directional change as well (Cheung, 2014; Heinze et al., 2018).

Path integration is a taxonomically widespread strategy likely to be used by animals living in habitats where visual landmark use is unreliable. Studies in desert ants demonstrated path integration using a sky compass and a stride integrator as an odometer (Wehner & Müller, 2006; Wittlinger et al., 2007), research in golden hamsters revealed the use of a single light source as a directional cue and self-motion information as an odometer for path integration (Etienne et al., 1990), investigations in stomatopods observed path integration underwater for the first time, albeit in a surface-bound animal (Patel & Cronin, 2020b), and studies in cichlid fish found initial evidence consistent with the use of path integration in aquatic vertebrates (Sibeaux et al., 2024). However, clear evidence for the use of path integration in fish is still lacking, and research in this group is neglected. Teleost fish represent half of all vertebrate species and are a highly diverse clade, but we have limited information on the diversity of their local navigational strategies (Volff, 2005). Additionally, fish exist in a three-dimensional medium and can move with six degrees of freedom, which may present additional challenges when navigating compared to surface-bound animals (Jeffery et al., 2013). Fish are therefore excellent models for navigation research, and studying path integration in fish could help determine how navigational strategies differ between animals which move in two-versus three-dimensional space.

An alternative to path integration is navigating by landmarks. This can range in complexity from simpler behaviours like beaconing, where an animal uses a landmark to identify the position of a goal, to more complex ones like piloting, where animals follow a sequence of landmarks to find a goal (Reese, 1989; Warburton, 1990). Landmarks can generally be classified as local, which directly indicate a specific location, or global, which can be used along with other cues to calculate a location (Braithwaite & Burt de Perera, 2006).

Many fish species live in coral reefs, which are landmark-rich environments. However, navigation by path integration could be beneficial for coral reef-dwelling fish in several situations. First, conditions with reduced visibility, such as increased water turbidity or plankton blooms, could make visual landmark use unreliable (Wenger & McCormick, 2013). Second, if landmarks become altered or unstable due to erosion, coral growth, or other disturbances, having additional navigational strategies available, like path integration, could be advantageous. Furthermore, while coral reefs are structurally complex overall, they may contain relatively featureless sections where landmark cues are scarce. In these cases, path integration could aid fish in finding their way back to their home range or shelter when other navigational cues are limited or disrupted. Evidence for path integration in coral reef fish would outline its use as a navigational strategy and provide insights into the evolution of this behavior and its underlying neural mechanisms.

We developed a novel experimental paradigm to test whether the coral reef-dwelling domino damsel fish (*Dascyllus trimaculatus*) could navigate back to a shelter location using path integration. Domino damsels are a very territorial species, with their mortality rate related to intraspecific competition for shelter (Holbrook & Schmitt, 2002). Shelter use is an important part of domino damsel life history, as shelters or rocky surfaces are used as a substrate to lay eggs. These fish tend to leave their shelter to forage and return to it for safety. We expected that it would be beneficial for domino damsels to path integrate, as it would allow them to return to safety rapidly and efficiently when encountering a predator or competitor. We therefore hypothesise that domino damsels can navigate back to a shelter location using path integration.

## Methods

### Overview

We tested whether domino damsel fish can navigate back to a shelter using path integration. We first allowed individuals to acclimatise to a circular experimental pool with two landmarks and a shelter overnight. The next morning, we placed a trap with food rewards in the centre of the pool. I recorded the fish as they swam from their shelter into the trap. Once trapped, we removed the landmarks and shelter from the pool. Fish were then released from the trap and free navigation movements were recorded and analysed.

### Animal Husbandry

We used 10 domino damsel fish (*Dascyllus trimaculatus*) in experiments sourced from a supplier (Tropical Marine Centre, Solesbridge Lane, Chorleywood, Hertfordshire, WD3 5SX). Individuals were wild caught and kept by the supplier for one month before being housed in the laboratory.

We housed individuals in separate 0.35 m × 0. 32 m × 0.60 m tanks within a single flow-through marine system. We enriched each tank with 1 cm depth of coral gravel substrate, a large rock, a cave shelter (13 cm × 12 cm × 8 cm Aqua One Marble Cave), and a 30 cm tall plastic plant. The fish were kept on a 12 h light/12 h dark cycle with fluorescent light, and fed individuals with crisp (TetraPRO Energy Multi-Crisps) in the morning and mysis shrimp in the afternoon to provide supplementary nutrients. We maintained water parameters at healthy levels for this species (Temperature: 26.0 °C, Salinity (specific gravity): 1.024, pH: 8.2, KH: 7-8 dKH, Nitrite: 0 ppm, Ammonia: 0 ppm, Nitrate: <20 ppm) by carrying out water tests, water changes, and tank cleans once per week. Room temperature, water temperature, and individual health were checked at each feed. We quarantined all individuals in this system for 4 weeks before starting experiments.

### Experimental Apparatus

The experiments were run in a 2 m-diameter blue plastic circular pool (Figure 2), which was filled with 10 cm depth coral gravel substrate and 20 cm depth water (from water surface to top of substrate). We kept pool water at identical parameters to the home tank system. To help maintain these parameters, we placed a water heater and a filter pump in the pool (Figure 2).

**Figure 2.**
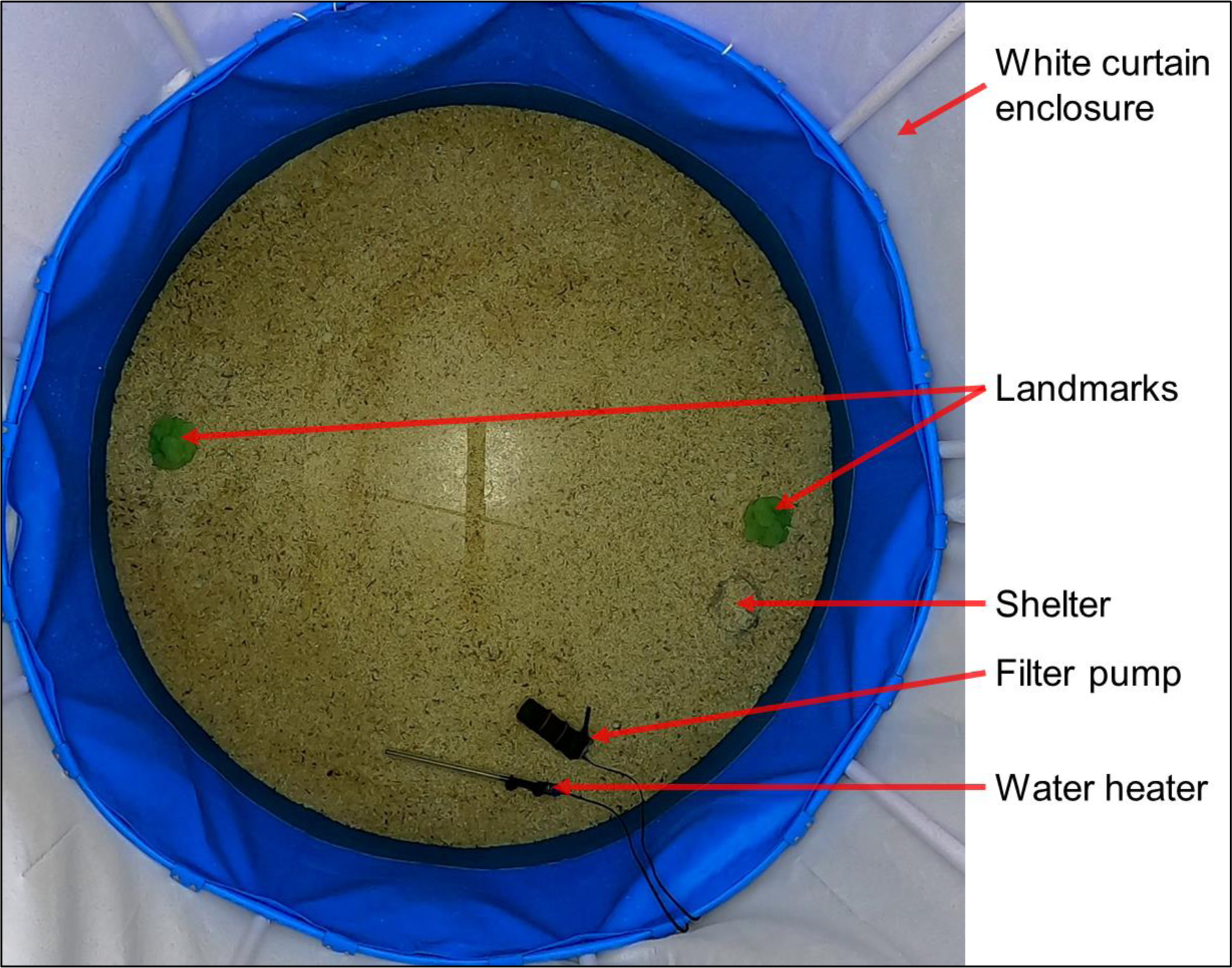
Picture taken on the GoPro showing the experimental pool prepared for fish acclimation.

During the acclimation period and in experiments, we placed two 14.0 cm × 14.6 cm × 18.1 cm polyurethane resin replica green brain coral landmarks and a shelter (14 cm × 11 cm × 8 cm Aqua One Marble Cave) in the pool. To reduce the shelter’s saliency, we covered it with coral gravel substrate using silicone. In this experiment, it was important for the shelter to be inconspicuous to prevent fish from remembering it as a salient landmark or beacon. In addition to covering the shelter with substrate, the shelter was slightly buried (i.e. positioned below the substrate level) forcing the fish to search for it when exploring the experimental pool. To reduce the likelihood of individuals using global visual cues from the laboratory room, we surrounded the pool with white curtains (Figure 2).

To record footage during acclimation and experiments, we positioned a GoPro Hero 8 Black 2 m above the centre of the pool (recording at 1440p, 30 fps, Linear FOV, see full view in Figure 2). To provide a low-latency live video feed of individual fish location, we positioned a USB webcam next to the GoPro and connected it to a laptop outside of the curtains. This second camera allowed me to monitor fish movement in real time for trapping.

### Acclimation

To allow fish to acclimatise to the pool overnight, we moved individuals from their home tank to the experimental pool in the afternoon the day before an experiment. we ensured that water conditions were identical before moving an individual into the pool to acclimatise by measuring temperature, salinity, and pH in the tank and pool. Once moved, we fed the fish and recorded the following hour of the acclimation period, allowing me to calculate an exploratory score for each individual.

### Experimental Setup

Before beginning an experimental trial, we measured pool water temperature. To eliminate confounding navigational cues within the pool, we removed the water heater and filter pump for the duration of the experimental trial. Stable external directional cues are essential for precise path integration in many species and the location of the sun, or even a single light source, can be used as a directional cue in animal path integration (Etienne et al., 1990; Heinze et al., 2018; Patel & Cronin, 2020b). To provide a directional cue analogous to the sun, we positioned a spotlight outside the experimental enclosure at a height of 128 cm and a horizontal distance 48 cm away from the pool, facing the curtains with a 20° negative inclination. This created a localised bright spot that the fish could use as a directional cue or heading indicator for path integration (Goodyear & Ferguson, 1969; Guilford & Taylor, 2014).

We then placed the trap into the centre of the pool (Figure 3a). The trap consisted of a 12.5 cm diameter by 25.3 cm height white opaque plastic cylinder with a 6.4 cm diameter entrance hole cut into one side 1.6 cm from the bottom. We used the trap to contain the fish during the trial and prevent it from directly observing any experimental manipulations. To get the trap to appear more like a natural shelter, we covered the exterior and interior with the coral gravel substrate. There was a 17.5 cm by 7.5 cm rectangular white opaque plastic door with attached steel weights behind the trap’s entrance hole, which opened or closed by sliding along plastic rails (Figure 3a). We were able to open, close, and remove the trap remotely by using fishing lines attached via a pulley system. To provide additional enrichment and encourage the fish to explore the central area of the pool, we placed a plastic plant by the door (Figure 3a). We then started each trial by setting the GoPro to record, adding mysis shrimp inside the trap, and remotely raising the door.

**Figure 3.**
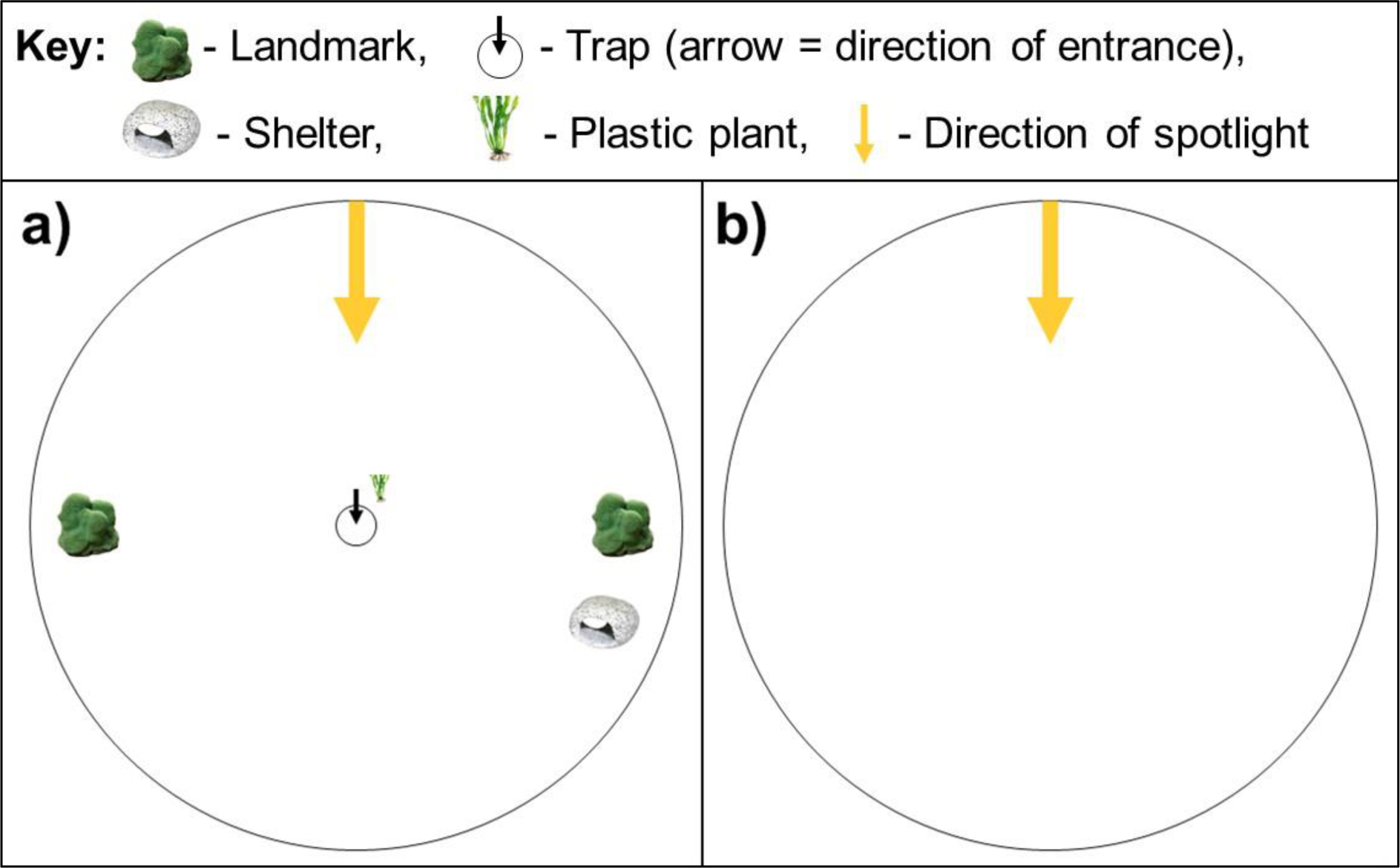
Layout of the pool at the start of an experimental trial (a), and layout of the pool after removing the shelter and landmarks and releasing the fish by vertically lifting out the trap (b).

### Experimental Procedure

We recorded fish movement in each trial for up to two hours, and ended experimental trials if individuals did not enter the trap in this period. As soon as we observed fish entering the trap on the live low-latency USB camera feed, we closed the trap remotely. We then removed the landmarks, the shelter, and the plastic plant (Figure 3b). To prevent fish from using chemical cues from the water or visual patterns in the substrate to identify the shelter location, we also mixed and smoothed out any disturbed substrate and water near the landmark and shelter locations. Finally, we released the fish by lifting the trap vertically out of the water using the pulley system. We recorded five minutes of fish free movement before ending the trial and used a large jug to return the fish to its home tank.

### Fish Tracking and Trajectory Analyses

Over 6 weeks of testing and 21 experimental trials attempted, only 6 of the 10 fish tested entered the trap. The following analyses have therefore been performed on the trajectories collected from these 6 individuals.

Fish movement in the acclimation and pre-trapping part of the experimental trial was tracked using AnimalTA (Chiara & Kim, 2023). Fish movement after release from the trap was recorded manually in MATLAB (MathWorks, 2023), and coordinates were extracted every second for one minute. We tracked post-release movement manually as the background subtraction tracking method used by AnimalTA was not suitable once the trap was removed from the water during a trial.

I focused on and analysed three navigational periods in detail:

1. Acclimation: To characterise fish exploratory behaviour during the acclimation period, we calculated average speed (cm/s), total distance travelled (cm), and area of the pool explored (cm^2^) in AnimalTA. we also recorded exploration latency (the first time at which each fish exited the shelter and became entirely visible on camera, s) and measured the greatest distance it swam from its shelter (cm). Each fish was given an exploratory index calculated by multiplying normalised versions of the above metrics.
2. Last outward trajectory: To characterise fish behaviour during the experiment before the fish was trapped, we analysed fish movement during the last outward trajectory. This is defined as the period between the fish leaving its shelter for the last time during the trial and it being trapped in the centre. We calculated the length (cm) and time taken (s) for the last outward trajectories in AnimalTA. To obtain measures of the tortuosity of the last outward trajectories, we calculated their straightness indices and sinuosities in R using the ‘trajr’ package (version 1.5.1; McLean & Skowron Volponi, 2018). The straightness index is defined as 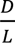, where *D* is the beeline distance between the first and last points in the trajectory and *L* is the path length travelled (Batschelet, 1981). Sinuosity, defined by Benhamou (2004), uses the mean step length, the mean cosine of turning angles, and the coefficient of variation of step length (standard deviation/mean) to calculate a measure of tortuosity.
3. Homeward path: To characterise fish behaviour after being released from the trap, we analysed manually recorded fish coordinates (extracted with MATLAB) using the ‘trajr’ package in R (McLean & Skowron Volponi, 2018; R Core Team, 2022). We used the above metrics of straightness index and sinuosity to determine the transition point between the homeward path ending and the initiation of search behaviour. We identified this transition through visual inspection of a graph of straightness index and sinuosity over time, and defined the end of the homeward path based on when two sequential coordinates showed a breakdown in straightness index or increase in sinuosity after a plateau (Figure 4).

**Figure 4.**
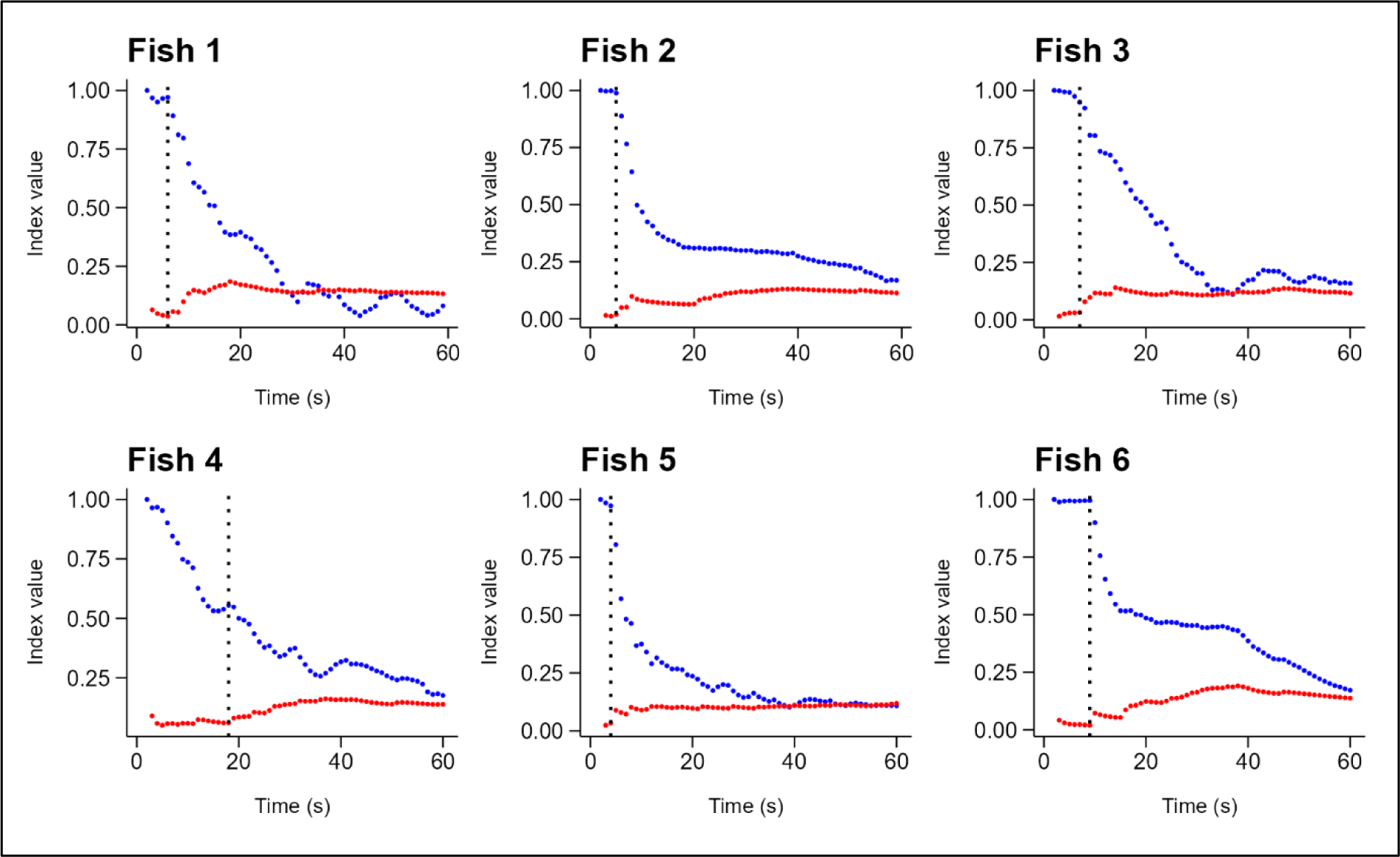
Straightness index (blue) and sinuosity (red) of the post-release path of each fish develop over time. The first shared large deviation in straightness index or sinuosity (dotted line) was used to define the end point of an individual’s homeward path.

Once the homeward paths were identified, we calculated the homeward path direction (angle between first and last homeward path coordinates, degrees), beeline distance (straight line distance between first and last homeward path coordinates, cm), length (cm), time (s), mean speed (cm/s), straightness index, and sinuosity. We then used the homeward path direction to calculate homeward direction error (absolute angular difference between homeward path direction and shelter direction, degrees). We also calculated the homeward beeline distance error (absolute difference between shelter distance and homeward beeline distance, cm) and the homeward path length error (absolute difference between shelter distance and homeward path length, cm).

### Statistical analysis

All statistics were carried out in R (version 4.2.2; R Core Team, 2022). We used circular statistics to analyse direction data, and the significance threshold α was set at 0.05.

#### 1 – Homeward path direction analysis

To investigate whether fish homeward path directions followed a uniform, unimodal, or bimodal distribution, we used a model-based approach with maximum likelihood with the R package ‘CircMLE’ (version 0.3.0; Fitak & Johnsen, 2017). The fish direction data were converted to circular class data using the package ‘circular’ (version 0.5.0; Agostinelli & Lund, 2023). These directions were then compared to ten direction models and the best fitted model was estimated using AIC values (for more details see Fitak & Johnsen, 2017). We also conducted a Rayleigh test of uniformity to determine if the distribution of homeward path directions differed significantly from a uniform distribution.

#### 2 – Homeward path direction relative to shelter

We next analysed whether homeward path directions differed significantly from direction to the shelter. First, we conducted a Watson’s two-sample test of homogeneity between the distribution of homeward path directions and a simulated distribution centred on the shelter location with the same concentration parameter as the homeward path directions. The concentration parameter used (3.095) was estimated during the maximum likelihood analysis with ‘CircMLE’ (Table S1).

Second, we calculated a circular 95% confidence interval around the mean homeward path direction to see if this contained the shelter direction. Mean direction was calculated using the R package ‘circular’ (version 0.5.0; Agostinelli & Lund, 2023), and confidence intervals were calculated by bootstrapping with replacement over 1000 iterations using the package ‘boot’ (version 1.3.28; Angelo Canty & B. D. Ripley, 2024).

#### 3 – Effect of last outward trajectory behaviour on homeward path behaviour

We next tested how homeward path behaviour was influenced by last outward trajectory behaviour by fitting a series of linear models using the R package ’lme4’ (Bates et al., 2015). We tested whether a selection of biologically relevant last outward trajectory metrics set as explanatory variables (last outward trajectory length, time, straightness index, and sinuosity) had any influence on homeward path behaviour (response variables included: homeward direction error, homeward beeline distance error, homeward path length error, time, mean speed, straightness index, and sinuosity). For all linear models fitted see Table S2.

#### 4 -Homeward path distance relative to shelter

We then analysed whether homeward path beeline distance or homeward path length differed significantly from distance to the shelter. We conducted a Welch Two Sample t-test between homeward path beeline distances and shelter distance. We also conducted a Wilcoxon rank sum exact test between homeward path lengths and shelter distance, as homeward path lengths were not normally distributed (Shapiro-Wilk normality test: W = 0.77291, p = 0.0331).

#### 5 -Effect of acclimation on homeward path behaviour

We then tested how homeward path behaviour was influenced by acclimation behaviour by fitting a series of linear models. We tested whether a selection of biologically relevant acclimation metrics set as explanatory variables (acclimation distance travelled, area explored, and exploratory index) had any influence on homeward path behaviour (response variables included: homeward direction error, homeward beeline distance error, homeward path length error, total time, mean speed, straightness index, and sinuosity). For all linear models fitted see Table S2.

#### 6 – Power analyses

We finally conducted post hoc power analyses to determine whether my experiment had an appropriate sample size to observe significant effects of exploratory and outward path behaviour on homing behaviour. We used the package ‘lme4’ to fit my models and determined the power of these models with the ‘*powerSim’* function in the R package ‘simr’ (Green & MacLeod, 2016). For the results of the power analyses, see the supplementary material.

## Results

### 1 – Homeward path directions followed a unimodal distribution

The directions fish swam on their homeward paths were best described by a unimodal distribution following a circular maximum likelihood approach (ΔAIC = 0, φ_1_ = 6.049, κ_1_ = 3.095, Table S1, Figure S1). The remaining circular distribution models fitted poorly described the homeward path direction data (ΔAIC ≥ 2.145, Table S1). We confirmed the non-uniformity of homeward path directions with a Rayleigh test (r = 0.8168, p = 0.0112). The homeward and search paths of each fish are shown in Figure 5a.

**Figure 5.**
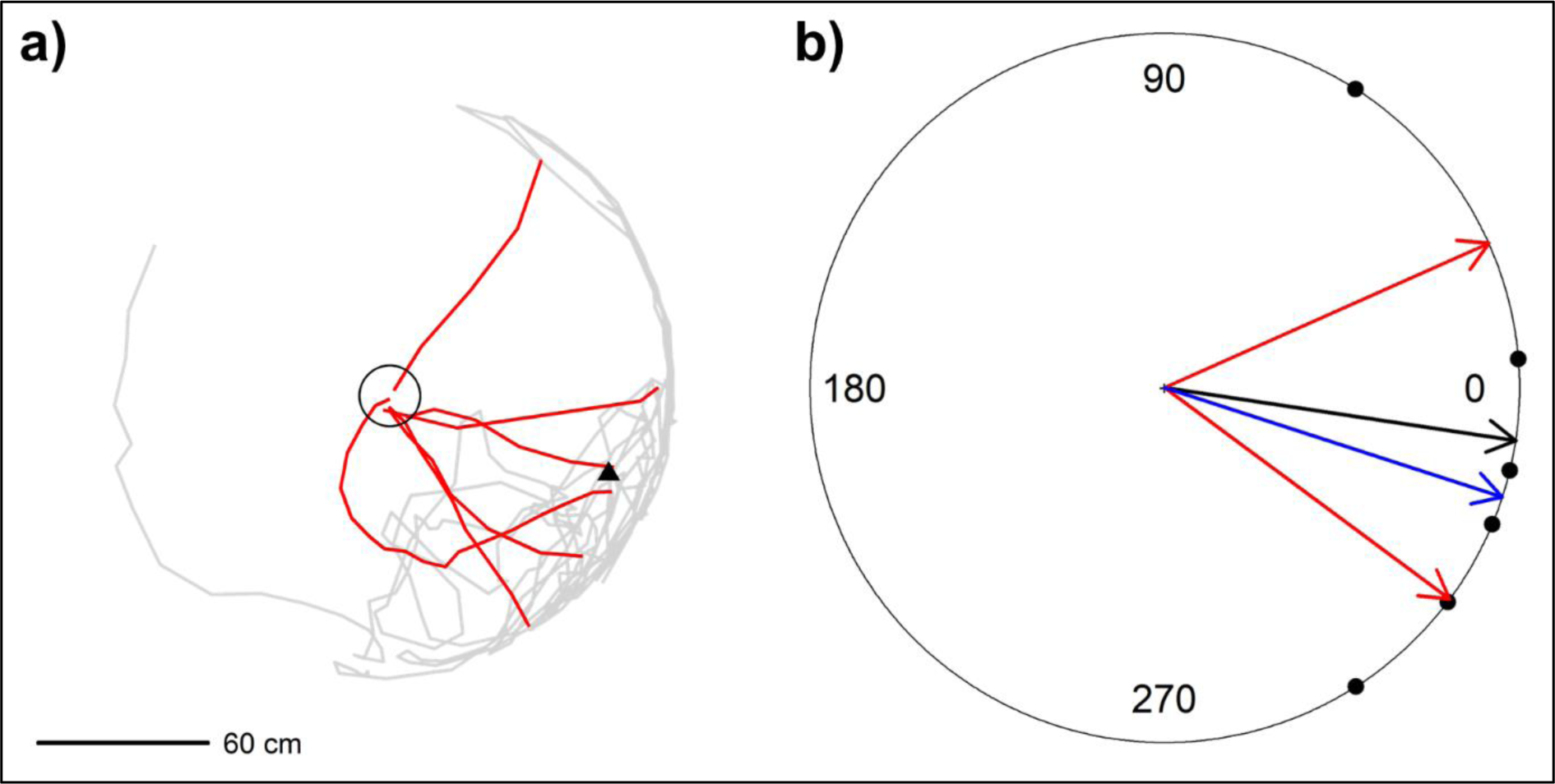
**(a)** Homeward (red) and search (grey) paths of each fish after release from the trap. Food trap location (circle) and previous shelter location (triangle) are also shown. **(b) Shelter angle is contained within the 95% confidence interval of mean homeward path direction.** Homeward path directions (black dots), mean homeward path direction (black arrow), 95% confidence interval of mean homeward path direction (red arrows), and shelter direction (blue arrow) are shown.

### 2 -Homeward path directions were oriented towards the shelter location

We did not find any significant difference between homeward path directions and shelter direction after conducting a Watson’s two-sample test of homogeneity between the homeward path directions and a simulated distribution centred on the shelter location with the same concentration parameter as the homeward path directions (Test statistic = 0.0486, Level 0.05 critical value = 0.187). Additionally, the direction of the shelter location was contained within a circular 95% bootstrap confidence interval for the mean homeward path direction (Figure 5b).

### 3 -Homeward direction error increased with last outward trajectory length

As hypothesised, homeward direction error increased with the length of the last outward trajectory. The linear model fitted between last outward trajectory length and homeward direction error explains a statistically significant and substantial proportion of variance (R^2^ = 0.74, F_(1, 4)_ = 11.42, p = 0.028, adjusted R^2^ = 0.60). Within this model the effect of last outward trajectory length is statistically significant and positive (β = 0.01, 95% CI [1.90 × 10^-3^, 0.02], t_(4)_ = 3.38, p = 0.028; Std. β = 0.86, 95% CI [0.15, 1.57], Figure 6). None of the other models fitted showed any significant effect of last outward trajectory behaviour on homeward path behaviour (Table S2).

**Figure 6.**
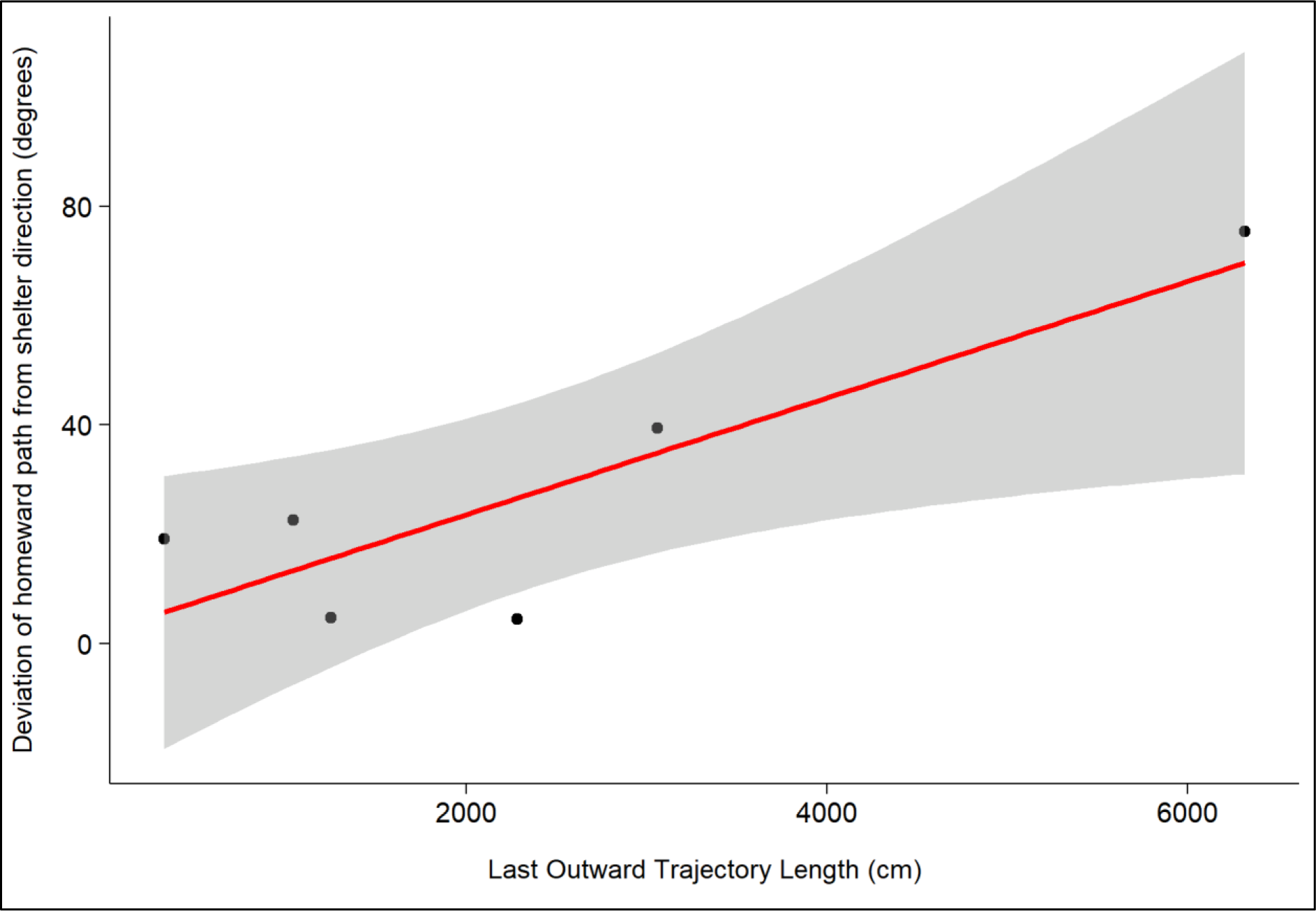
Homeward path direction error increases with last outward trajectory length.

### 4 -Homeward path lengths and beeline distances were significantly greater than shelter distance

Both homeward beeline distance and homeward path length were significantly greater than the distance to the shelter. The Welch Two Sample t-test suggested that beeline length is significantly greater than shelter distance (mean of beeline length = 88.17, mean of shelter distance = 78.00, difference = 10.17, 95% CI [1.51, 18.83], t(6.05) = 2.87, p = 0.028; Cohen’s d = 1.66, 95% CI [0.16, 3.07]). Additionally, the Wilcoxon rank sum exact test result suggested that homeward path length is significantly greater than beeline length (W = 32.00, p = 0.026; r (rank biserial) = 0.78, 95% CI [0.34, 0.94]).

### 5 -Acclimation behaviour did not significantly influence homeward path behaviour

No significant effects of any of the biologically relevant acclimation explanatory variables (acclimation distance travelled, area explored, and exploratory index) were observed on any of the homeward path response variables (homeward path length error, homeward beeline distance error, total time, mean speed, straightness index, and sinuosity). Results for all of the linear models fitted can be found in Table S2.

## Discussion

The results of this study provide evidence that the coral reef-dwelling domino damselfish (*Dascyllus trimaculatus*) can exhibit homing behaviour consistent with the use of path integration. After being released from a central location with no visual landmarks or their previous shelter present, the fish swam in a unimodally clustered distribution of directions that was not significantly different from the actual direction towards their previous shelter. This suggests the fish were attempting to navigate directly back to their shelter location rather than orienting randomly.

The error in the directional accuracy of the fish’s homeward paths increased with the length of the last outward trajectory taken before being released from the central trap location. This relationship between outward path length and homing error fits the expectations of path integration and has been observed in many species, where errors in an internally calculated estimation of directional change and distance covered accumulate over larger travel distances due to noise (Cheung, 2014; Heinze et al., 2018; Merkle et al., 2006). Noise is present at all physical scales and impacts every level of the nervous system. Its manifestations include fundamental quantum uncertainty, stochasticity of neural spikes, and environmental perturbation, distortion, or obstruction of sensory cues (Faisal et al., 2008). In path integration, noise can take the form of random environmental disturbances or alterations of path integration inputs, computations, or outputs within the sensorimotor system. These cannot be predicted or detected by the animal so result in error around a home vector. While exploratory behaviour during acclimation did not significantly influence homing accuracy, effects may emerge with larger sample sizes (see power analyses in the supplementary material).

Many other navigational strategies can be ruled out or seem unlikely based on the experimental conditions used and the results obtained. The use of local landmark-based navigational strategies like piloting and beaconing is highly unlikely due to the removal of the shelter and landmarks as visual cues. Additionally, the curtain surrounding the pool would limit the use of global landmarks outside the arena, except for the provided spotlight. The lack of influence acclimation behaviour appears to have on homeward movement, combined with the relatively homogeneous nature of the pool environment, casts doubt on the possibility that fish navigate to their shelter by learning a map based on cues in their environment (for example, based on the geometric relationships of objects present). The use of olfactory navigation is also unlikely due to the deliberate mixing of water and substrate to disrupt olfactory cues. While the fish could theoretically use magnetoreception, sound, or electroreceptive cues for homing, there is no direct evidence supporting reliance on these strategies. Though it is possible that geomagnetic cues could be used as a compass cue for path integration, the use of a magnetic coordinate map to navigate is unlikely due to the small spatial scale of the experimental pool and the nature of the homing behaviour observed. The use of hydrodynamic cues to navigate is also improbable due to the still water conditions in the pool. As homing error increases with longer outward path lengths, path integration emerges as the most viable and parsimonious explanation for the observed homing behaviours in this experimental setting.

Path integration is a robust navigational strategy, allowing an animal to return to the starting point of a journey without using familiar positional cues. It works through continuously estimating position relative to a reference point using only directional and distance cues derived from an animal’s own locomotion. This enables an animal to efficiently return home along the most direct route at any place or moment on its journey, even in featureless or unfamiliar terrain (Etienne & Jeffery, 2004; Müller & Wehner, 1988b). Path integration can become more reliable when combined with other strategies, like using landmarks or environmental cues to calibrate a homing vector (Etienne et al., 1996; Patel & Cronin, 2020a). Due to its ability to be used with other navigational strategies and its presence in disparate animal taxa, it is unsurprising that path integration might be one of many navigational strategies employed by fish.

Despite the fact that animals can path integrate using purely self-generated (or idiothetic) directional cues, it is unlikely that this was the case for the individuals in this experiment. The fact that idiothetic rotational estimates must be used cumulatively, together with the impact of noise on these estimates, means idiothetic path integration leads to rapid accumulation of error around the animal’s direction estimate. In contrast, the use of a stable external (or allothetic) directional cue allows the constant calibration of an animal’s direction estimate relative to this cue, so directional error does not accumulate. Irrespective of the type of directional cue used, errors around an animal’s estimation of distance travelled always accumulate linearly with increasing outward path length (Cheung et al., 2007; Heinze et al., 2018). Due to the lack of significant impact of last outward trajectory straightness or sinuosity on homeward direction error, together with the significant linear increase of homeward direction error with increasing last outward trajectory length, the use of stable external directional cues appears more likely than the use of self-generated idiothetic directional cues in this experiment.

Both the spotlight provided during experimental trials and the Earth’s magnetic field could have served as directional cues for fish path integration in this experiment. Mosquitofish and rodents have been shown to use point light sources or sun-analogous cues to estimate their direction (Etienne et al., 1990; Goodyear & Ferguson, 1969). Therefore, the spotlight provided in this experiment may have functioned similarly as a global landmark or heading indicator that individuals could use as a directional cue for path integration (Guilford & Taylor, 2014). Additionally, the use of magnetoreception is widespread among fish, so individuals could have used geomagnetic cues as a compass for path integration (Naisbett-Jones & Lohmann, 2022). The use of other compass or directional cues, like the position of stars, skylight polarisation, a self-generated cognitive map, or odour cues, was highly unlikely due to the nature of the experimental conditions and behaviour observed.

As well as estimating direction, path integration also requires an animal to estimate distance travelled. Fish could use a wide variety of cues to estimate this, including optic flow, proprioception, energy expenditure, vestibular cues, or water flow. Recent research has demonstrated that fish can estimate their distance travelled using optic flow (Karlsson et al., 2022; Sibeaux et al., 2022). While other mechanisms like proprioception, energy expenditure, and vestibular cues can be used to estimate distance travelled in terrestrial animals, they are yet to be explored in fish (Jacob et al., 2017; Proffitt et al., 2003; Wittlinger et al., 2007). Lastly, although the use of water flow cues to estimate distance travelled may be plausible in fish, evidence supporting this behaviour is lacking.

While providing novel evidence supporting the use of path integration in coral reef fish, there are limitations to the current study that could be addressed through future research. First, the specific compass and odometer cues used for path integration in fish remain unidentified. Systematically manipulating potential external directional cues, like shifting the position of the artificial light source or altering the local magnetic field, could determine which directional cues are prioritised. Additionally, altering potential odometer cues, like changing the visual characteristics of the pool environment or altering the flow of water, could discern how these fish estimate distance travelled.

Second, future research could better characterise how error accumulates around the estimated length of the home vector. While we found that homeward path length and homeward beeline distance both differed significantly from shelter distance, it’s possible that fish homing behaviour was influenced by the relatively close proximity of the pool wall. Individuals may have associated the pool wall with their shelter location, so may have continued toward the wall having followed a homing vector. Similar behaviour has been observed in hamsters: when startled in an experiment, individuals followed a homing direction towards the arena edge, then searched for their nest entrance by following the arena border (Siegrist et al., 2003). Furthermore, any stress associated with being trapped in the centre or the absence of their shelter may have induced a wall-following behavioural response (thigmotaxis), leading to fish continuing on toward the wall after following their homing vector (Champagne et al., 2010; Sharma et al., 2009). Future experiments in a larger arena could position the shelter further from any walls to both limit any influence of walls on homing, and to more definitively characterise path integration behaviour in fish. For example, this would allow better investigation into how homeward path length and beeline distance compare to shelter distance, and if errors in these distance measures relative to shelter distance also accumulates linearly with outward path length. This could also enable the characterisation of systematic search behaviour in fish without the influence of a nearby wall.

Third, experiments introducing controlled outward paths with predetermined lengths and tortuosities, instead of variably-terminating last outward trajectories, could more precisely characterise how homing error accumulates with increasing outward path length and directional change. Encouraging fish to swim through a series of controlled outward paths of predetermined lengths could more precisely characterise how path integration error accumulates with increasing outward path length. Additionally, allowing fish to swim through controlled outward paths of differing tortuosities could identify whether fish use stable external directional cues or self-generated cues to estimate direction when path integrating. However, restricting the movement of fish using controlled paths could lead to biased results. Individuals might follow the boundaries of any controlled paths, and the necessary recording of other stimuli (like the position of path boundaries) might distract from direction or distance monitoring.

Furthermore, fish exist in a three-dimensional medium and can move with six degrees of freedom, but this experiment only analyses movement in two dimensions. Future research could build on my initial findings that fish follow homing trajectories consistent with path integration and use an experimental arena with greater three-dimensional complexity to explore how this navigational behaviour changes with greater use of an additional spatial dimension. Lastly, increased sample sizes would likely provide greater statistical power to detect any weaker effects of acclimation or outward path behaviour on homing behaviour (see power analysis in the supplementary materials). By building upon this foundation with targeted experiments, we can gain deeper knowledge into the specific sensorimotor mechanisms underpinning path integration in coral reef fish.

Despite these limitations, our results expand our understanding of path integration abilities to include coral reef fish. Path integration has previously been demonstrated in terrestrial invertebrates like desert ants, terrestrial vertebrates like rodents, and marine arthropods like stomatopod crustaceans (Etienne et al., 1990; Müller & Wehner, 1988b; Patel & Cronin, 2020b). The finding that a coral reef fish possesses similar path integration capabilities implies this navigational strategy either evolved early in animals or potentially multiple times through convergent evolution across disparate taxa.

From an applied perspective, understanding the sensory cues and mechanisms behind fish path integration could aid in predicting patterns of fish movement and distribution, which is relevant for ecology, conservation, and fisheries management (Boström et al., 2011). Theoretically, details on how path integration functions in fish could help reveal the evolutionary origins of this ability and shed light on the neural processes governing spatial navigation systems.

In summary, this study provides novel evidence that the coral-reef dwelling domino damsel fish *Dascyllus trimaculatus* possess the ability to follow a homing trajectory consistent with path integration over spatial scales relevant to their ecology and life history. Future research could investigate the specific odometer and compass cues these fish use for path integration. Overall, the results add coral reef fish to the list of taxa employing this widespread navigation strategy.

## Acknowledgements

We thank Christine Soper for her help with animal husbandry and all the technicians from the John Krebs Field station for helping with the fish. We thank Dr Cait Newport for lending us the experimental pool and spotlight and all Fish Lab people for their feedback on the project. We thank the Human Frontier Science Program and the Department of Biology of the University of Oxford for funding.

## Supplementary Materials

### Supplementary Tables and Figures

**Table S1.** Output from CircMLE. The output from the ‘circ_mle’ function for all 10 models of orientation using default parameters. Homeward path directions were used as the input data.

| Model | M2A | M2B | M2C | M3B | M4A | M1 | M4B | M3A | M5B | M5A |
| --- | --- | --- | --- | --- | --- | --- | --- | --- | --- | --- |
| Params | 2 | 2 | 3 | 3 | 5 | 3 | 4 | 0 | 4 | 2 |
| $\varphi_1$ | 6.049 | 5.858 | 6.052 | 5.939 | 5.853 | 2.73 | 5.984 | NA | 1.816 | 2.832 |
| $\kappa_1$ | 3.095 | 8.738 | 2.999 | 5.381 | 5.495 | 2.526 | 4.472 | 0 | 2.426 | 2.72 |
| $\lambda$ | 1 | 0.5 | 0.749 | 0.5 | 0.749 | 0.251 | 0.749 | 1 | 0.251 | 0.5 |
| $\varphi_2$ | NA | NA | NA | 9.081 | 0.723 | 5.872 | 9.125 | NA | 6.092 | 5.974 |
| $\kappa_2$ | 0 | 0 | 0 | 0 | 5.152 | 2.526 | 0.475 | 0 | 2.426 | 2.72 |
| Likelihood | 5.835 | 6.908 | 6.573 | 7.102 | 5.122 | 7.679 | 6.715 | 11.027 | 7.094 | 9.805 |
| Convergence | 0 | 0 | 0 | 0 | 0 | 0 | 0 | 0 | 0 | 0 |
| AIC | 15.67 | 17.815 | 19.146 | 20.203 | 20.243 | 21.358 | 21.43 | 22.055 | 22.189 | 23.61 |
| AICc | 19.67 | 21.815 | 31.146 | 32.203 | Inf | 33.358 | 61.43 | 22.055 | 62.189 | 27.61 |
| BIC | 15.253 | 17.399 | 18.521 | 19.579 | 19.202 | 20.734 | 20.597 | 22.055 | 21.356 | 23.193 |
| ΔAIC | 0 | 2.145 | 3.476 | 4.534 | 4.573 | 5.689 | 5.761 | 6.385 | 6.519 | 7.94 |
| ΔAICc | 0 | 2.145 | 11.476 | 12.534 | Inf | 13.689 | 41.761 | 2.385 | 42.519 | 7.94 |
| ΔBIC | 0 | 2.145 | 3.268 | 4.325 | 3.949 | 5.48 | 5.344 | 6.801 | 6.103 | 7.94 |
| Relative Likelihoods | 1 | 0.342 | 0.176 | 0.104 | 0.102 | 0.058 | 0.056 | 0.041 | 0.038 | 0.019 |
| AIC Weights | 0.5165 | 0.17672 | 0.09085 | 0.05354 | 0.05248 | 0.03005 | 0.0289 | 0.02122 | 0.01984 | 0.0097 |
| Evidence Ratio | NA | 2.923 | 5.686 | 9.649 | 9.842 | 17.19 | 17.82 | 24.348 | 26.038 | 52.987 |

**Table S2.** Summary of results for all linear models tested.

| Response variable | Explanatory variable(s) | R<sup>2</sup> | Adj. R<sup>2</sup> | p-value |
| --- | --- | --- | --- | --- |
| Homeward direction error (degrees) | Acclimation distance travelled (cm) | 0.09093 | -0.13630 | 0.56140 |
|  | Acclimation area explored (cm<sup>2</sup>) | 0.19630 | -0.00462 | 0.37890 |
|  | Exploratory index | 0.19860 | -0.00172 | 0.37580 |
|  | Last outward trajectory length (cm) | 0.74060 | 0.67580 | **0.02780\*** |
|  | Last outward trajectory time (s) | 0.64820 | 0.56020 | 0.05328 |
|  | Last outward trajectory straightness index | 0.12050 | -0.09940 | 0.50020 |
|  | Last outward trajectory sinuosity | 0.00180 | -0.24770 | 0.93630 |
| Homeward beeline distance error (cm) | Acclimation distance travelled (cm) | 0.34940 | 0.18670 | 0.21660 |
|  | Acclimation area explored (cm<sup>2</sup>) | 0.29240 | 0.11550 | 0.26790 |
|  | Exploratory index | 0.38180 | 0.22720 | 0.19110 |
|  | Last outward trajectory length (cm) | 0.32070 | 0.15090 | 0.24100 |
|  | Last outward trajectory time (s) | 0.34600 | 0.18260 | 0.21900 |
|  | Last outward trajectory straightness index | 0.06634 | -0.16710 | 0.62220 |
|  | Last outward trajectory sinuosity | 0.01156 | -0.23440 | 0.83900 |
| Homeward path length error (cm) | Acclimation distance travelled (cm) | 0.21570 | 0.01959 | 0.35300 |
|  | Acclimation area explored (cm<sup>2</sup>) | 0.47950 | 0.34930 | 0.12730 |
|  | Exploratory index | 0.46530 | 0.33160 | 0.13550 |
|  | Last outward trajectory length (cm) | 0.02824 | -0.21470 | 0.75000 |
|  | Last outward trajectory time (s) | 0.00411 | -0.24490 | 0.90400 |
|  | Last outward trajectory straightness index | 0.00391 | -0.24510 | 0.90600 |
|  | Last outward trajectory sinuosity | 0.16590 | -0.04267 | 0.42300 |
| Homeward time (s) | Acclimation distance travelled (cm) | 0.12680 | -0.09146 | 0.48840 |
|  | Acclimation area explored (cm<sup>2</sup>) | 0.16130 | -0.04844 | 0.43000 |
|  | Exploratory index | 0.26760 | 0.08444 | 0.29330 |
|  | Last outward trajectory length (cm) | 0.07201 | -0.16000 | 0.60710 |
|  | Last outward trajectory time (s) | 0.00534 | -0.24330 | 0.89060 |
|  | Last outward trajectory straightness index | 0.00018 | -0.24980 | 0.97990 |
|  | Last outward trajectory sinuosity | 0.04756 | -0.19060 | 0.67800 |
| Homeward mean speed (cm/s) | Acclimation distance travelled (cm) | 0.00080 | -0.24900 | 0.95750 |
|  | Acclimation area explored (cm<sup>2</sup>) | 0.00754 | -0.24060 | 0.87010 |
|  | Exploratory index | 0.00622 | -0.24220 | 0.88190 |
|  | Last outward trajectory length (cm) | 0.03717 | -0.20350 | 0.71440 |
|  | Last outward trajectory time (s) | 0.00073 | -0.24910 | 0.95940 |
|  | Last outward trajectory straightness index | 0.00643 | -0.24200 | 0.88000 |
|  | Last outward trajectory sinuosity | 0.00303 | -0.24620 | 0.91800 |
| Homeward path straightness index | Acclimation distance travelled (cm) | 0.08362 | -0.14550 | 0.57830 |
|  | Acclimation area explored (cm<sup>2</sup>) | 0.27620 | 0.09522 | 0.24830 |
|  | Exploratory index | 0.23990 | 0.04990 | 0.32400 |
|  | Last outward trajectory length (cm) | 0.10420 | -0.11980 | 0.53260 |
|  | Last outward trajectory time (s) | 0.05641 | -0.17950 | 0.65040 |
|  | Last outward trajectory straightness index | 0.00122 | -0.24850 | 0.94765 |
|  | Last outward trajectory sinuosity | 0.15110 | -0.06110 | 0.44627 |

| Response variable | Explanatory variable | R<sup>2</sup> | Adj. R<sup>2</sup> | p-value |
| --- | --- | --- | --- | --- |
| Homeward path sinuosity | Acclimation distance travelled (cm) | 0.05111 | -0.18610 | 0.66670 |
|  | Acclimation area explored (cm<sup>2</sup>) | 0.04035 | -0.19960 | 0.70270 |
|  | Exploratory index | 0.00089 | -0.24890 | 0.95530 |
|  | Last outward trajectory length (cm) | 0.04330 | -0.19590 | 0.69240 |
|  | Last outward trajectory time (s) | 0.05590 | -0.18010 | 0.65190 |
|  | Last outward trajectory straightness index | 0.00004 | -0.24990 | 0.99040 |
|  | Last outward trajectory sinuosity | 0.02820 | -0.21470 | 0.75000 |

**Figure S1.**
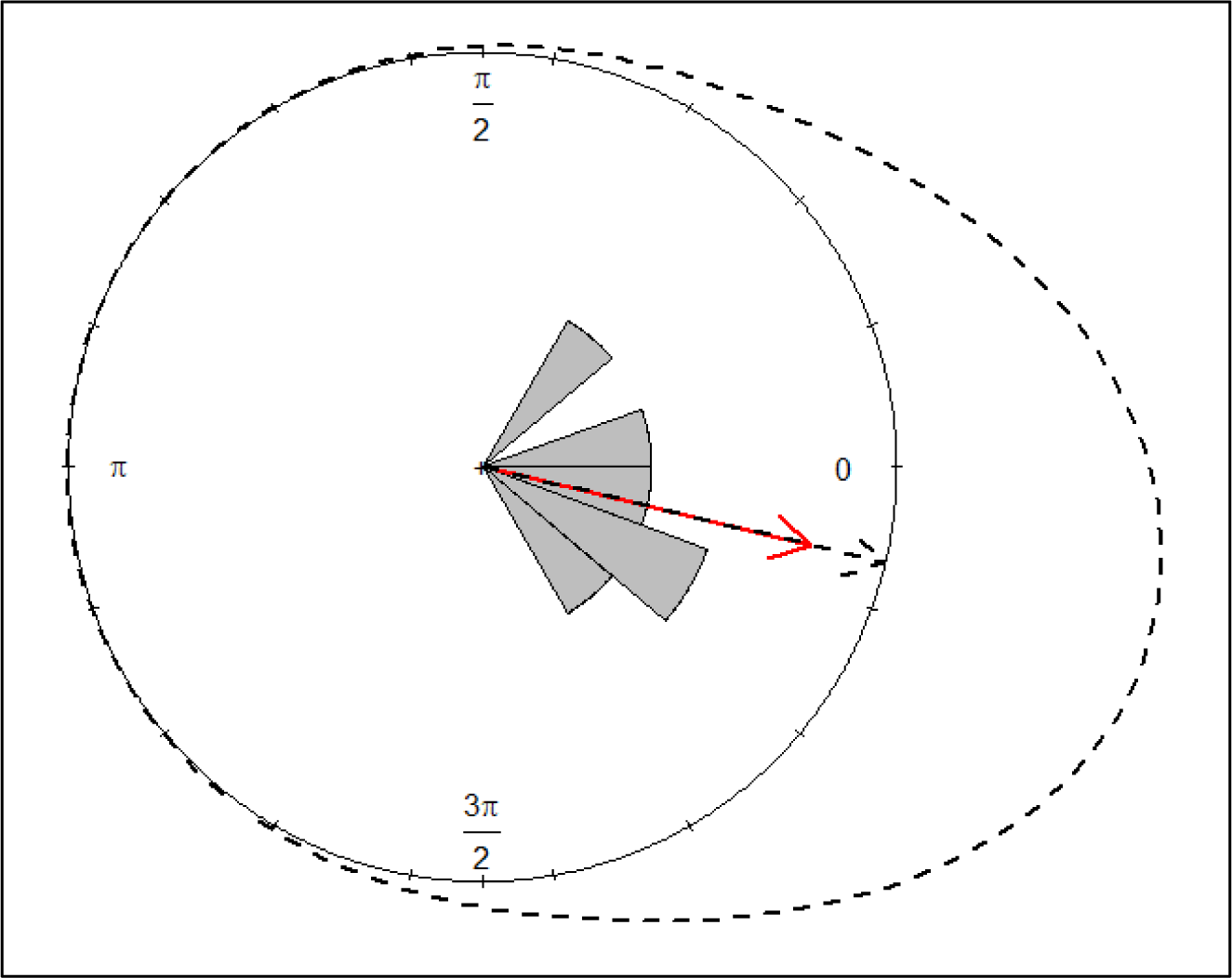
Best fit model for homeward path directions. Circular histogram (grey bars) and mean vector (red arrow) of homeward path directions. The density (dotted line) and mean angles (dotted arrows) for the best fit model, model M2A, are shown. The plot was produced using the ‘plot_circMLE’ function in the CircMLE package in R.

### Linear model power analyses

#### Linear Models

We conducted post hoc power analyses to determine whether our experiment was adequately powered to observe significant effects of exploratory and outward path behaviour on homing behaviour. We chose to analyse the power of four models in particular:

1. *homeward direction error ∼ last outward trajectory length*. We analysed the power of this model to determine whether our experiment had an appropriate sample size to detect whether homeward direction error accumulates with outward path length. Error accumulation is expected to occur if path integration is used to navigate.
2. *homeward direction error ∼ last outward trajectory straightness*. A significant effect of last outward trajectory tortuosity (here measured as straightness index) on homeward direction error could imply that individuals are using self-generated idiothetic directional cues rather than stable external directional cues when path integrating. We therefore analysed the power of this model and estimated an appropriate sample size to detect any effect.
3. *homeward direction error ∼ last outward trajectory sinuosity*. Similarly to (2), a significant effect in this model could imply individuals are using idiothetic cues, as sinuosity is another measure of path tortuosity.
4. *homeward direction error ∼ exploratory index*. A significant effect of exploratory index on homeward directional error would imply that fish could learn some aspects of the experimental pool before trials and use these learned cues to navigate. We therefore estimated an appropriate sample size to detect this effect.

#### *Model 1:* homeward direction error ∼ last outward trajectory length

We fit a linear model between last outward trajectory length and homeward direction error using the package 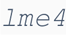, and determined the power of this model with the 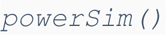 function in the 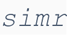 package:

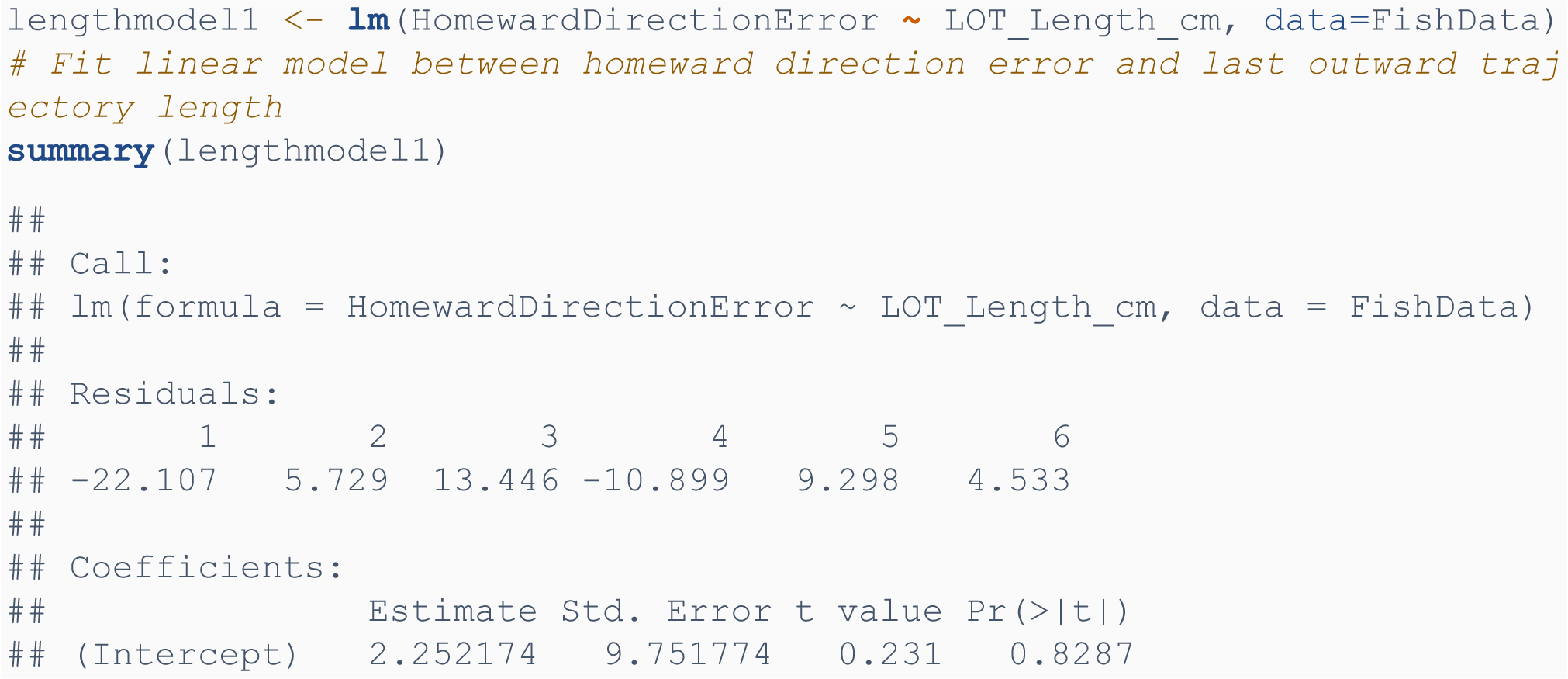
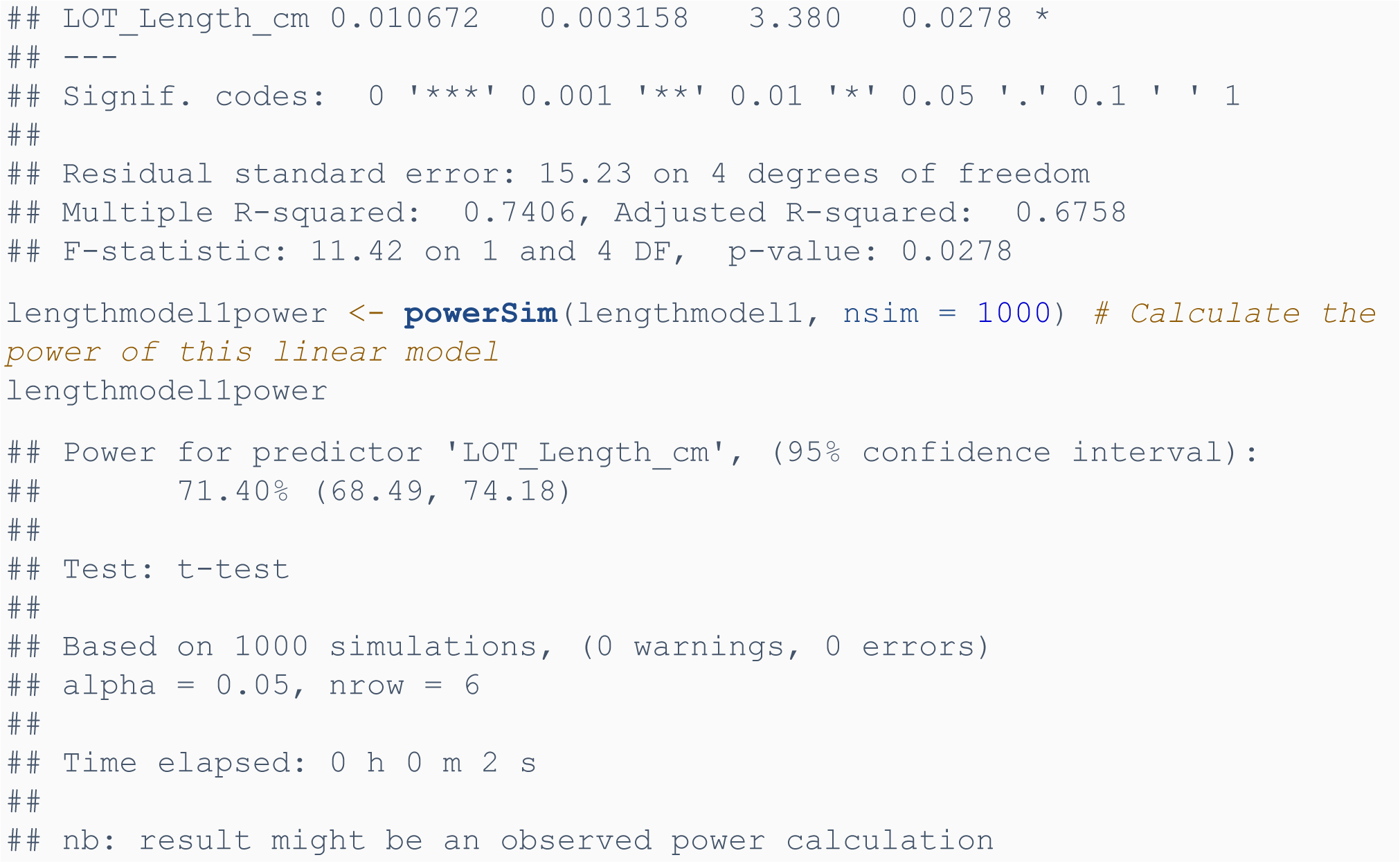

Due to the power observed (71.40%) being less than the generally accepted level of 80%, we extended the model to 10 fish to estimate the power at this sample size:

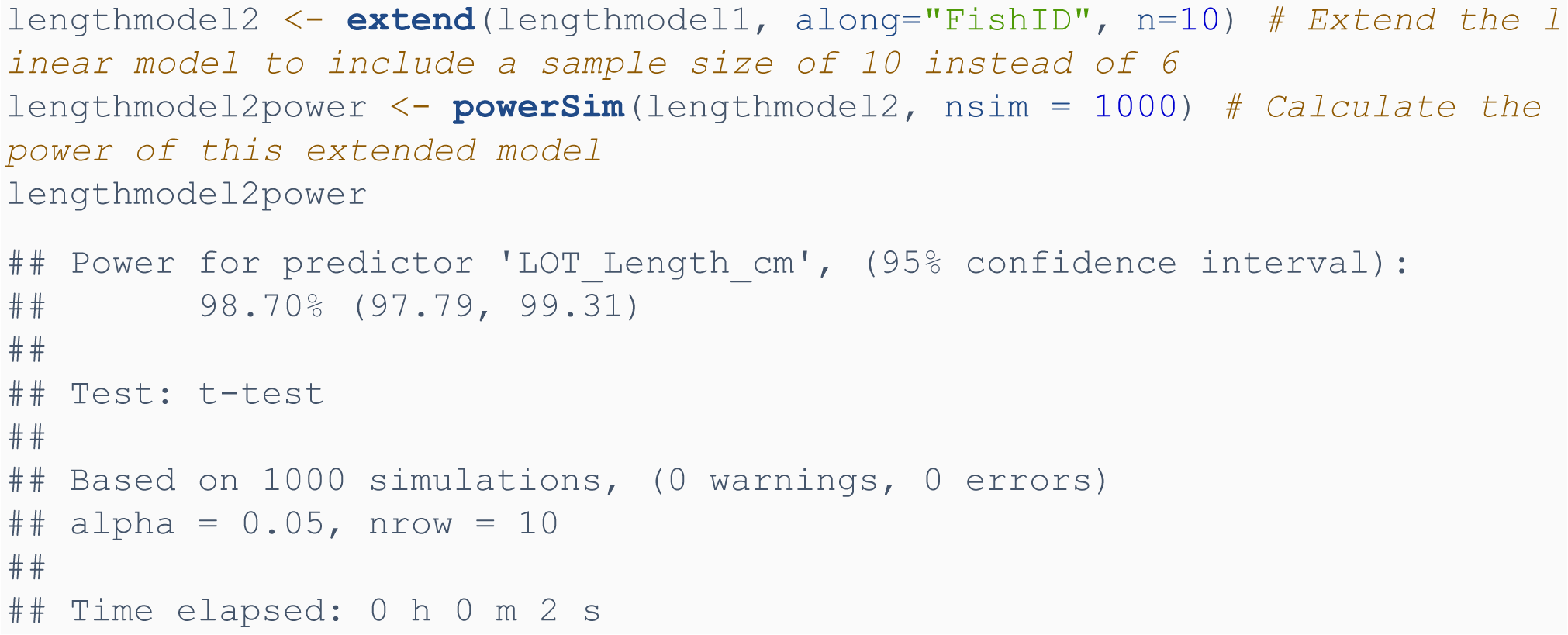

We then plotted a power curve to determine the minimum appropriate sample size for this model to have a power greater than 80%:

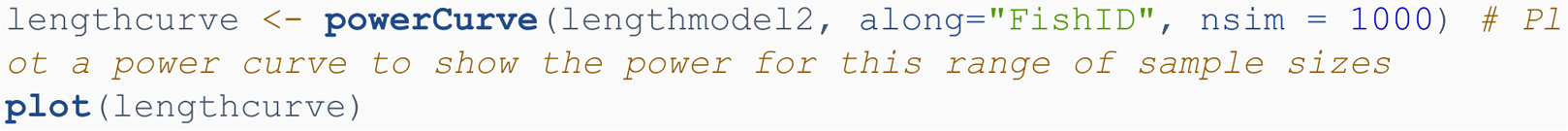

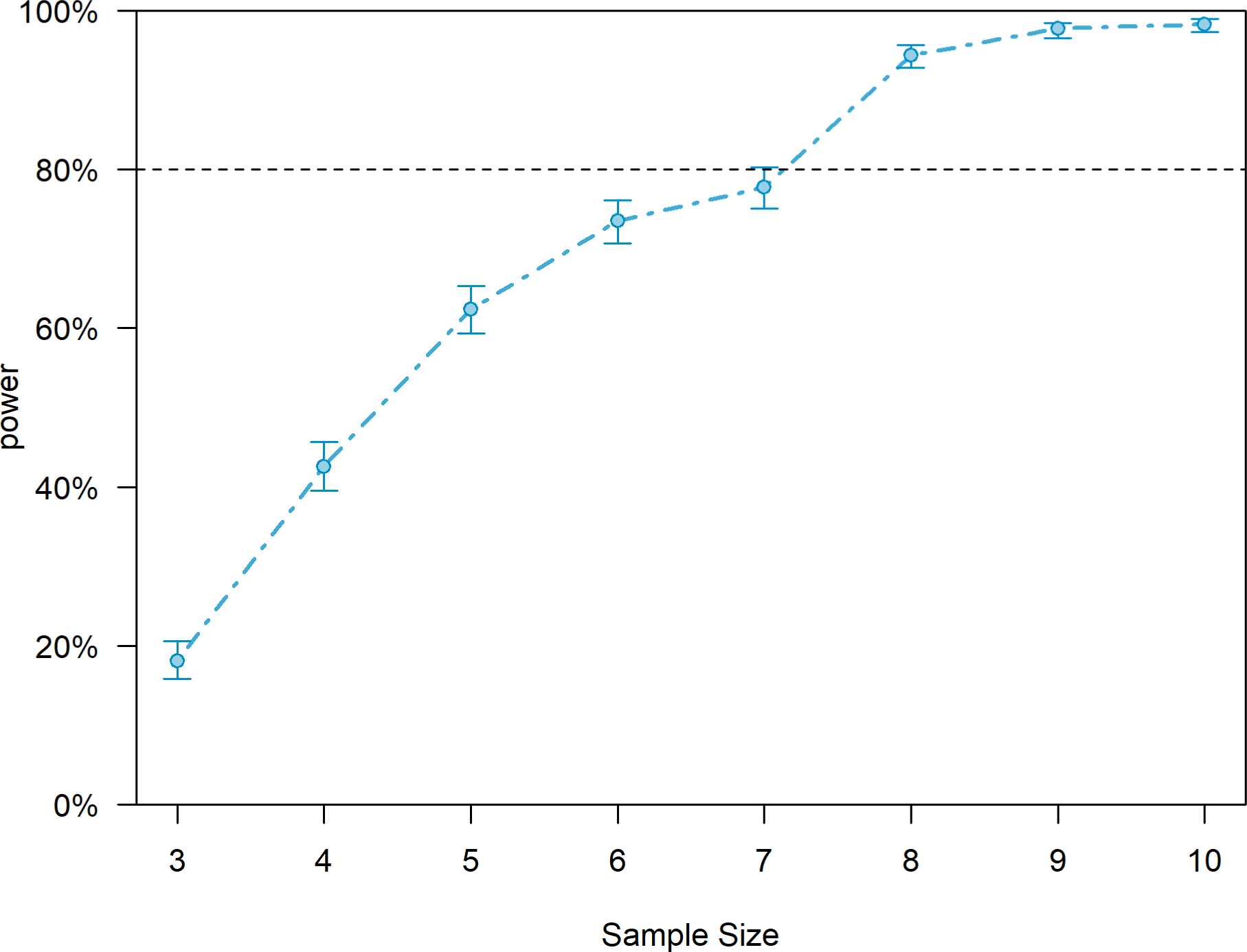

The resulting power curve suggests future experiments should use at least 8 fish to detect a significant effect of last outward trajectory length on homeward direction error.

#### *Model 2:* homeward direction error ∼ last outward trajectory straightne*ss*

We fit a linear model between last outward trajectory straightness and homeward direction error and determined the power of this model:

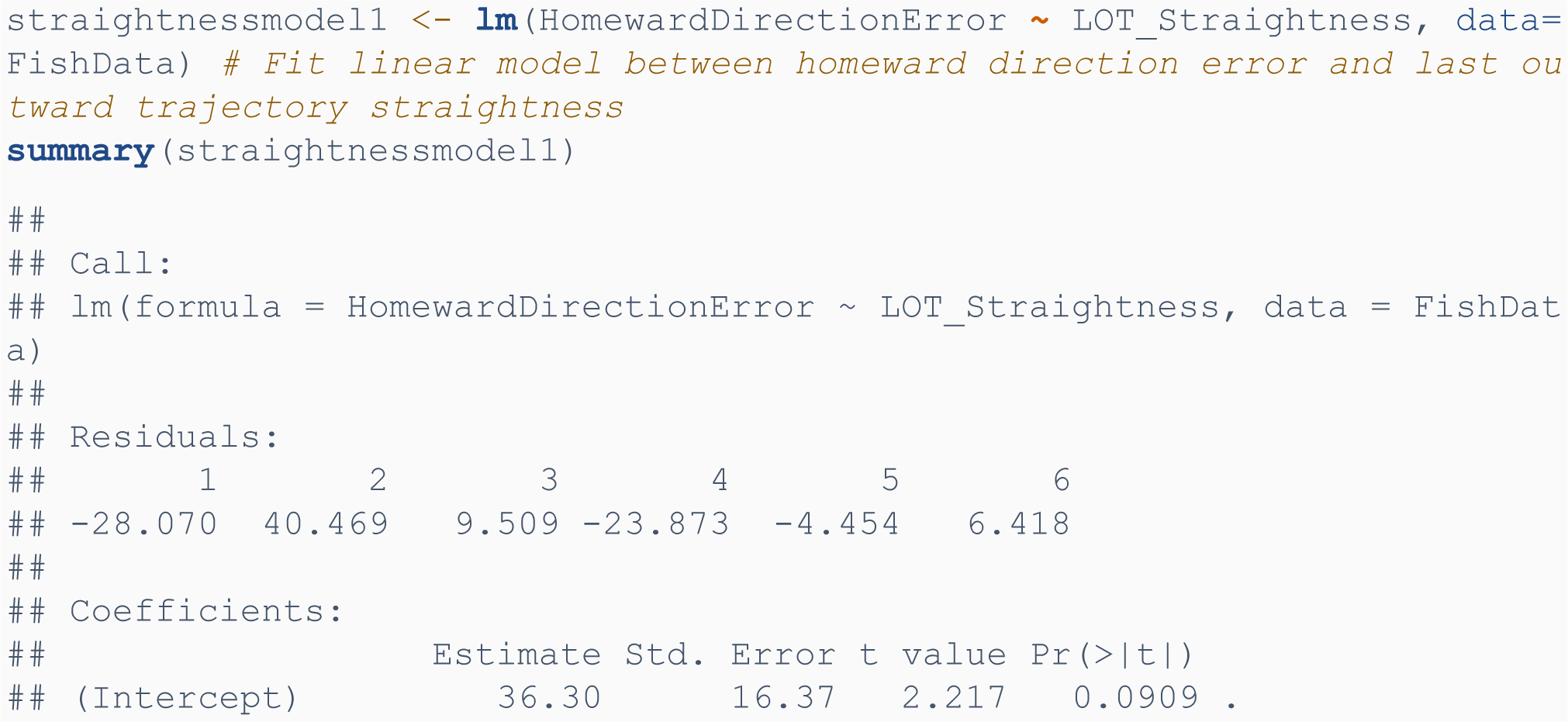
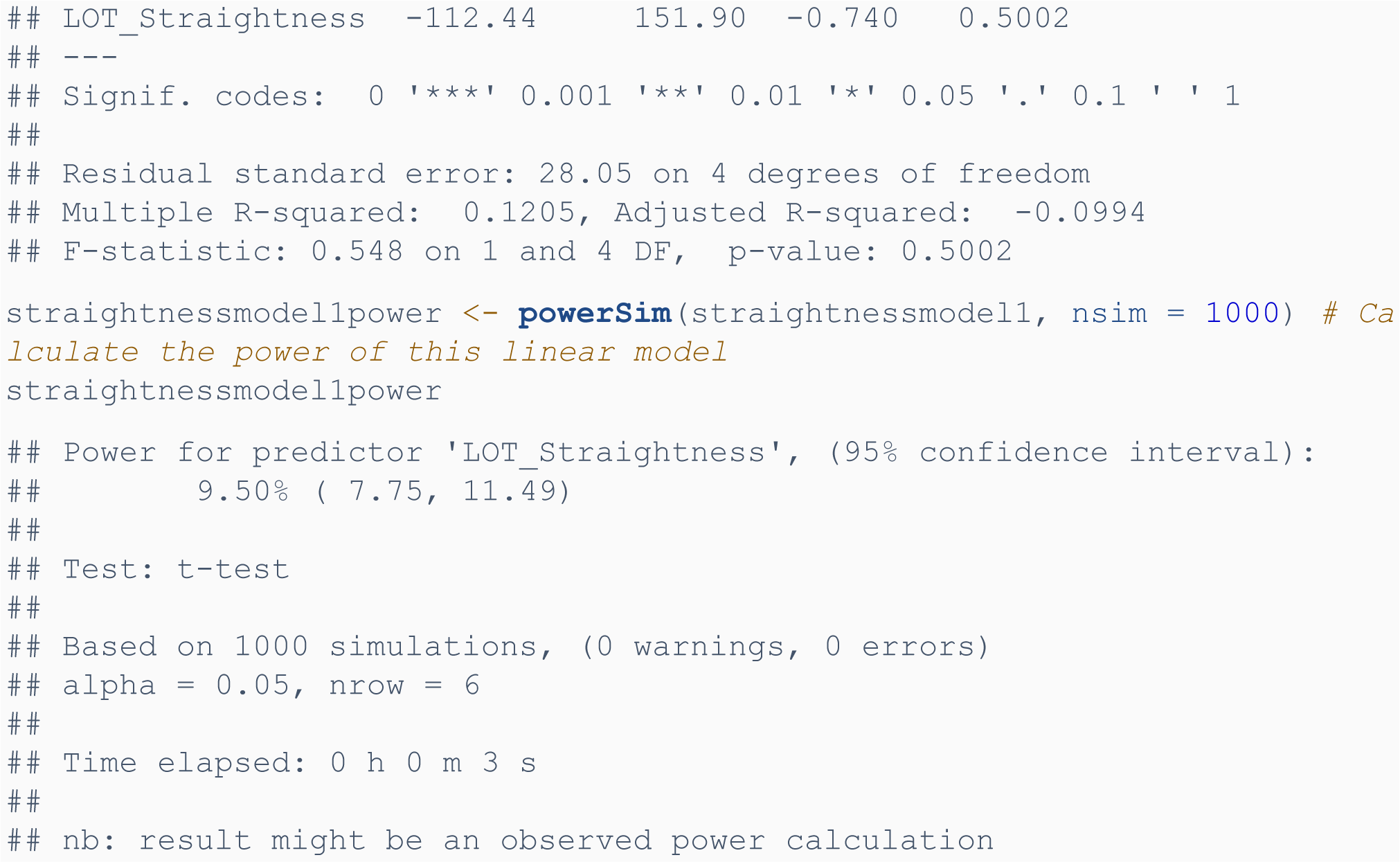

Due to power observed (9.50%) being much less than the generally accepted level of 80%, We extended the model to 100 fish to estimate the power at this sample size:

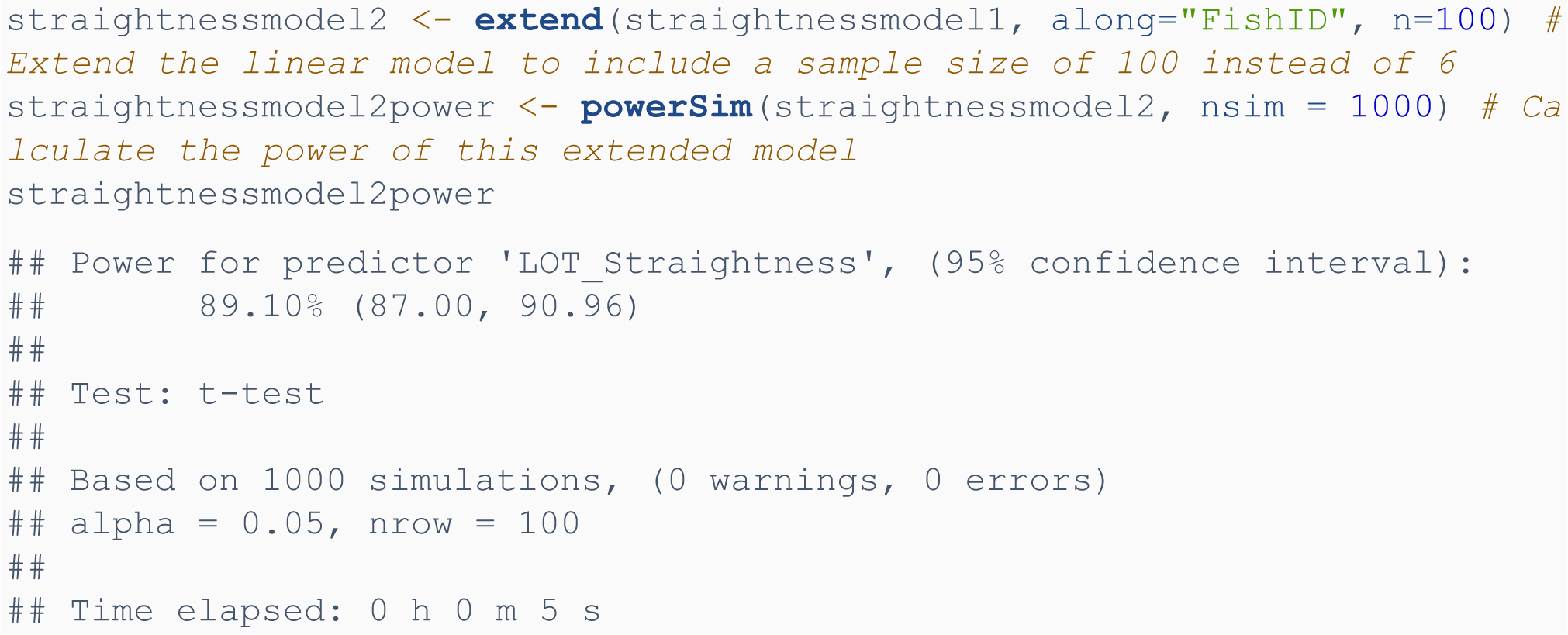

We then plotted a power curve to determine the minimum appropriate sample size for this model to have a power greater than 80%:

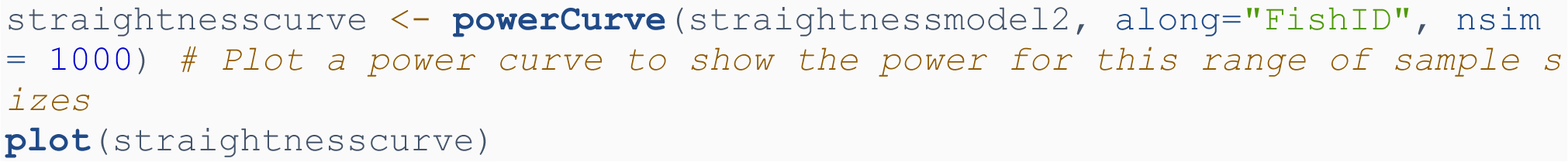

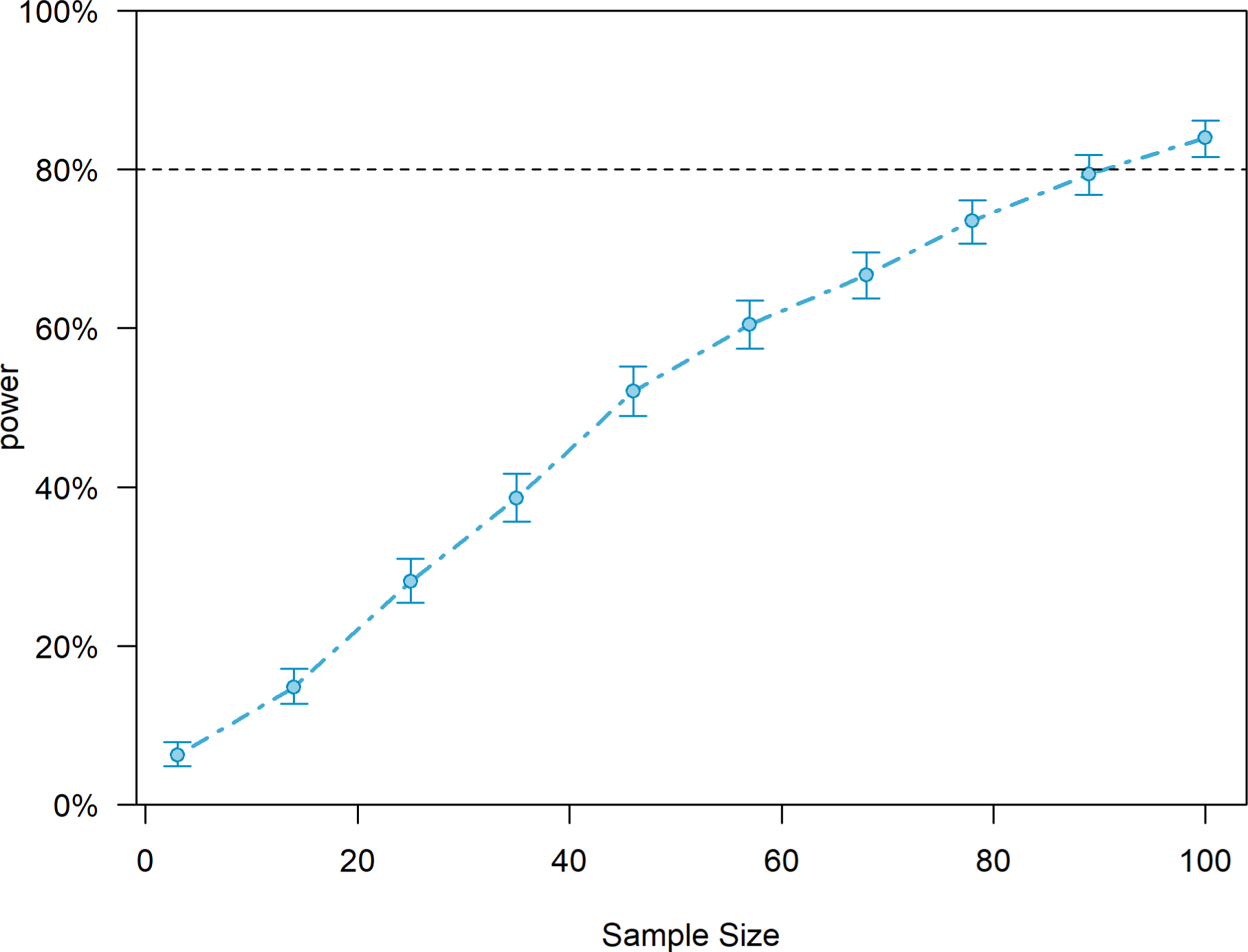

The resulting power curve suggests future experiments should at least 90 fish to detect a significant effect of last outward trajectory straightness on homeward direction error.

#### *Model 3:* homeward direction error ∼ last outward trajectory sinuosity

We fit a linear model between last outward trajectory sinuosity and homeward direction error and determined the power of this model:

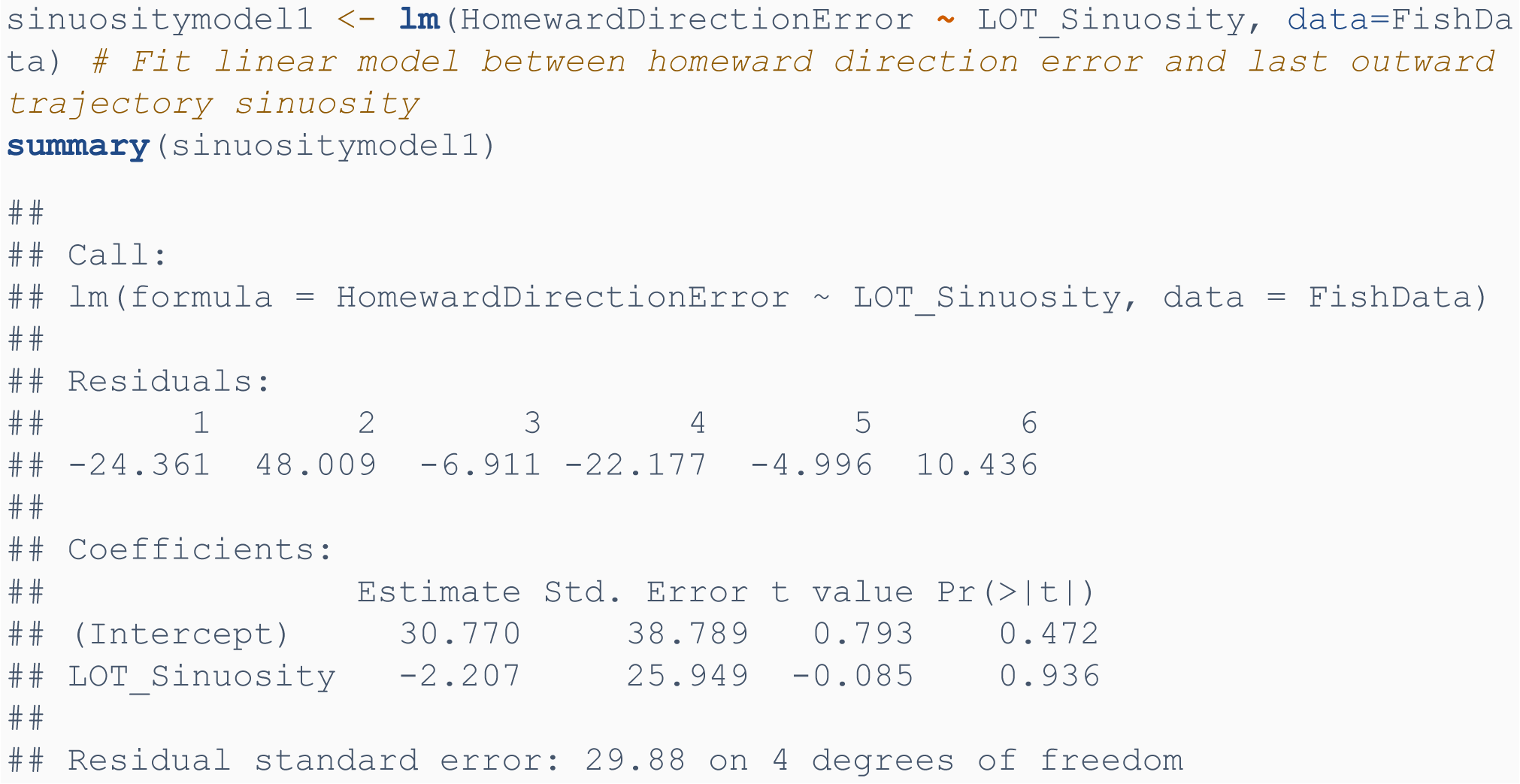
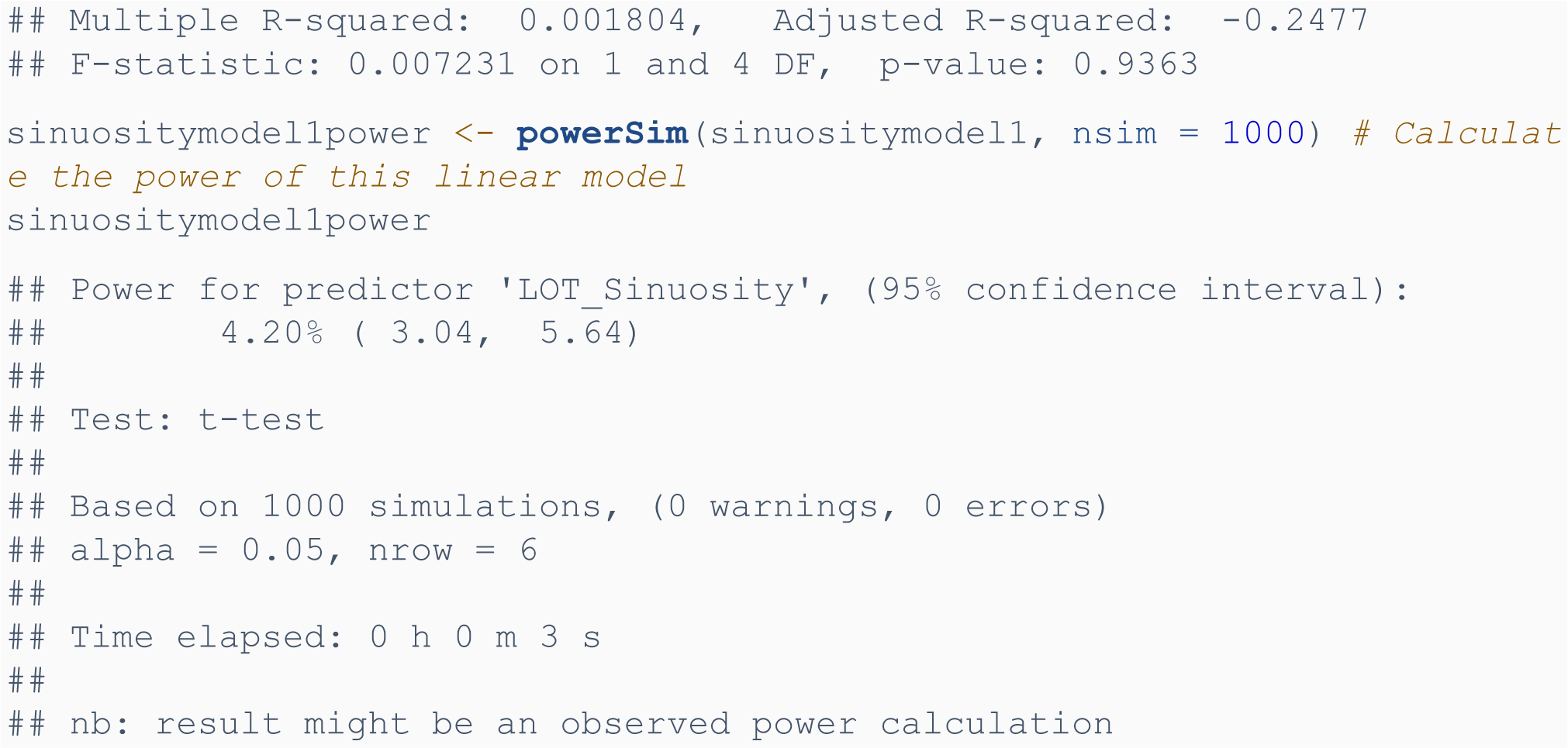

Due to power observed (4.20%) being much less than the generally accepted level of 80%, we extended the model to 6,000 fish to estimate the power at this sample size:

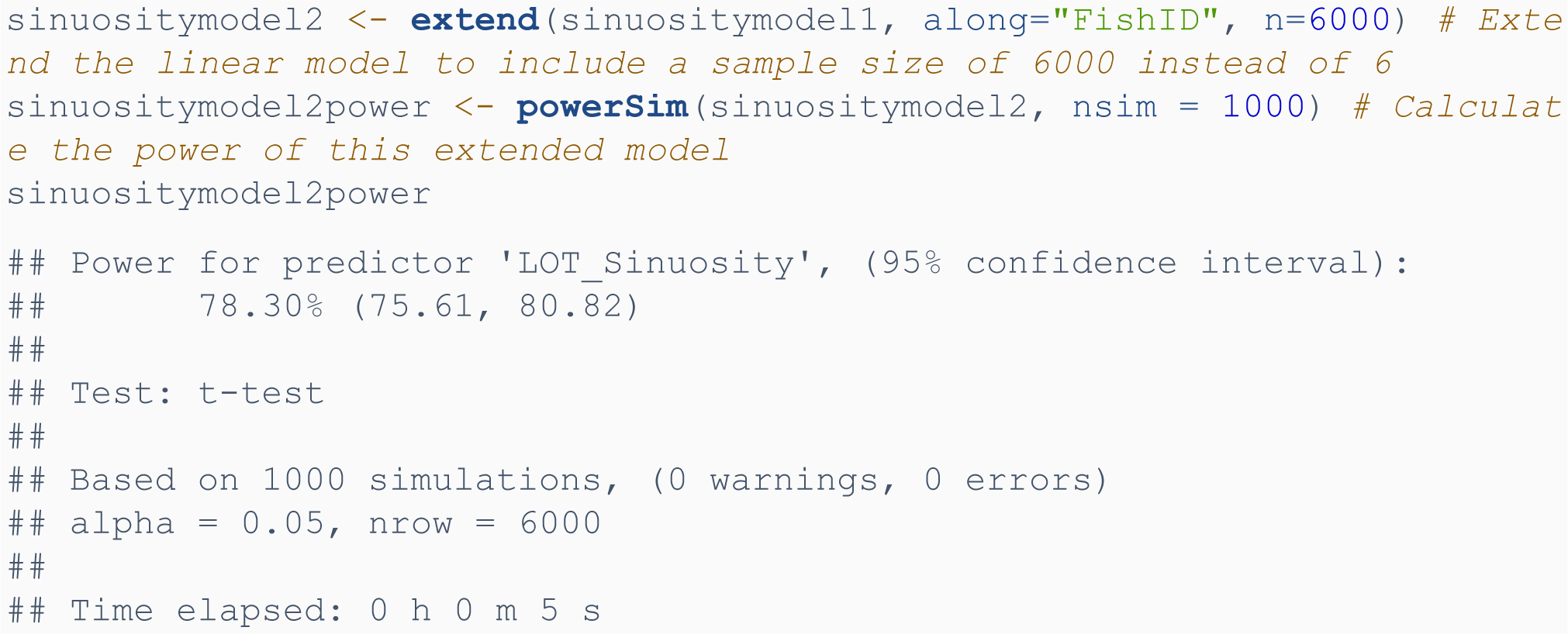

As the power of the model only appears to approach 80% when using a massive sample size (at least 6,000 fish), future research aiming to determine whether fish use idiothetic directional cues or stable external directional cues to path integrate should use a different experimental design that tests use of idiothetic vs allothetic directional cues more directly.

#### *Model 4:* homeward direction error ∼ exploratory index

We fit a linear model between exploratory index and homeward direction error and determined the power of this model:

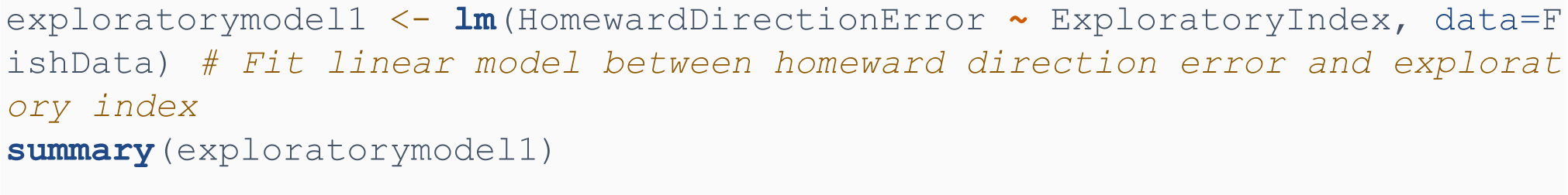
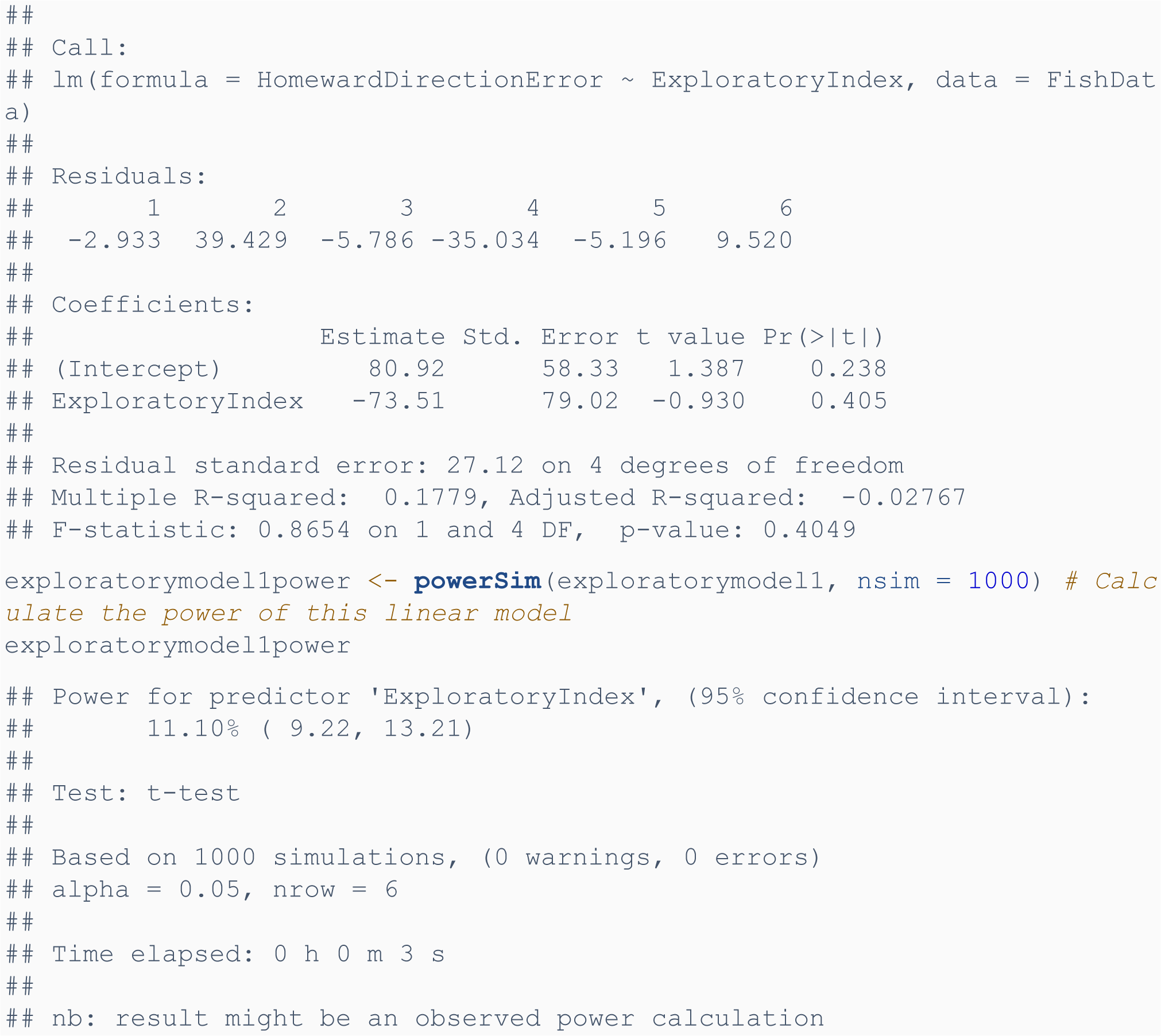

Due to power observed (11.10%) being much less than the generally accepted level of 80%, we extended the model to 60 fish to estimate the power at this sample size:

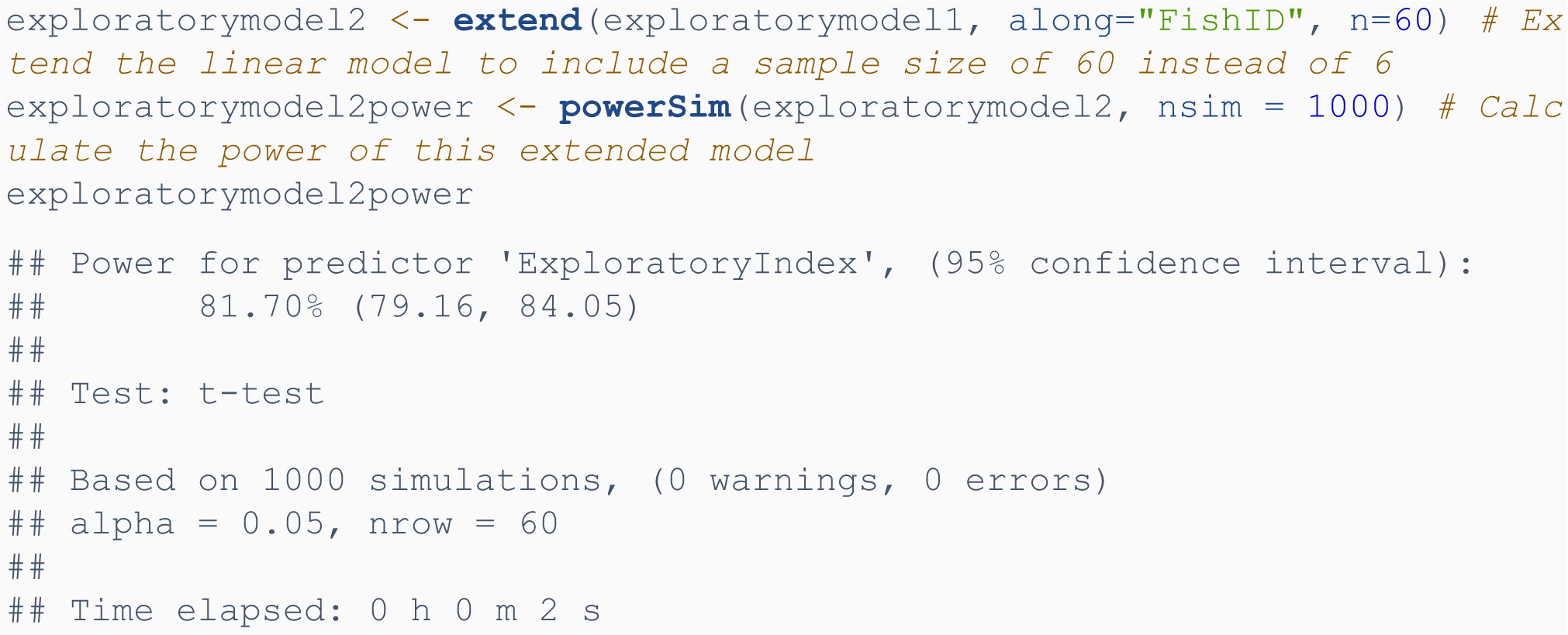

We then plotted a power curve to determine the minimum appropriate sample size for this model to have a power greater than 80%:

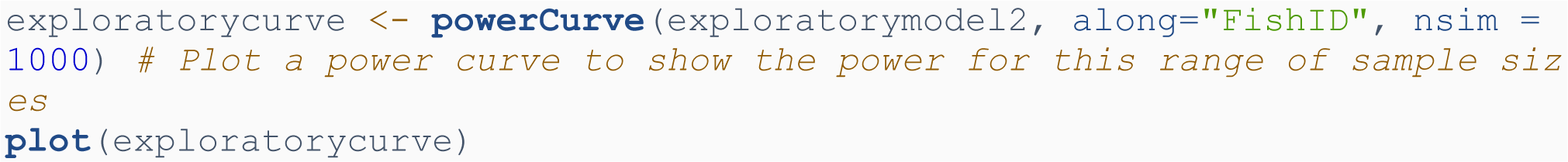

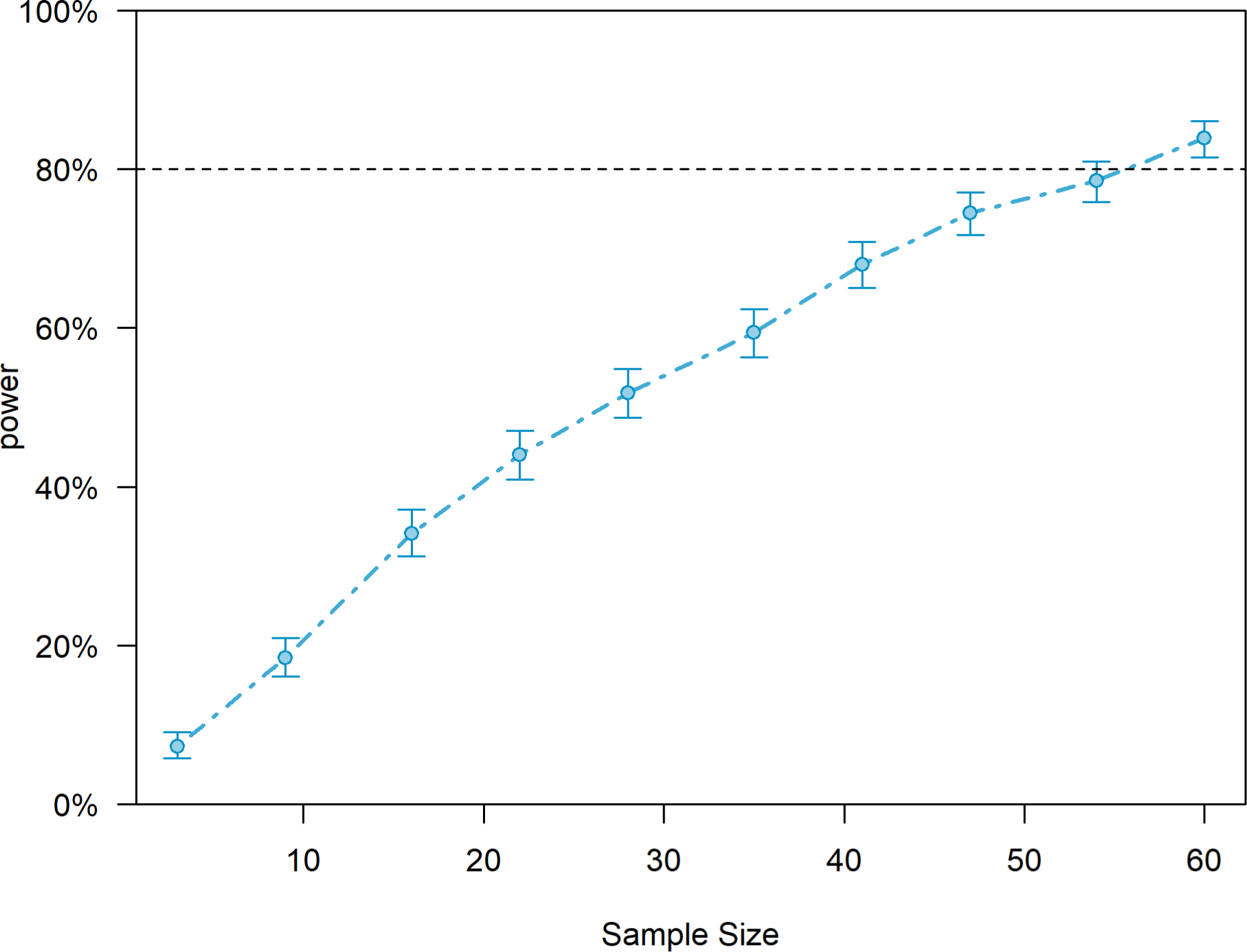

The resulting power curve suggests future experiments should include at least 60 fish to detect a significant effect of exploratory behaviour on homeward direction error.

### Conclusions

Although slightly underpowered, this experiment used a somewhat appropriate sample size to detect whether homeward directional error accumulated with increasing outward path length. The experiment was only slightly underpowered, with a power of 71% instead of the generally accepted 80%. Larger sample sizes or different experimental designs are needed to be able to more conclusively determine whether homeward path behaviour is influenced by other outward path behaviour or by exploratory behaviour in the acclimation period.

## Notes

### Competing Interest Statement

The authors have declared no competing interest.

### Summary of Updates

We corrected typos in the manuscript

